# Core MCC of *Arabidopsis thaliana*: structural dissection provides new insights into Mad2, BubR1 and CDC20

**DOI:** 10.64898/2026.11.06.764920

**Authors:** Eduardo Alves Gamosa de Oliveira, Emanuel Victor Nogueira Gotardo, Paula Guimarães Custodio, Gabriela Pimentel Serodio, Izabela Silva dos Santos, Marco Antonio Lopes Cruz

**Affiliations:** Laboratório de Biotecnologia Vegetal, Instituto de Biodiversidade e Sustentabilidade, Centro de Ciências da Saúde, Universidade Federal do Rio de Janeiro (BIOTEC/NUPEM/CCS/UFRJ) - Av. Amaro Reinaldo dos Santos Silva, 764, São José do Barreto, Macaé. CEP 27965-045; Department of Mechanistic Cell Biology, Max Planck Institute of Molecular Physiology (MPI) - Otto-Hahn-Strasse 11, 44227 Dortmund, Germany; Departamento de Bioquímica, Instituto de Química, Universidade de São Paulo Avenida Professor Lineu Prestes, 748, São Paulo SP 05508-000, Brasil

## Abstract

The spindle assembly checkpoint (SAC) is a mechanism executed mainly by the mitotic checkpoint complex (MCC) to monitor the anchoring of mitotic spindle microtubules to ensure correct chromosome segregation. Although knowledge of protein‒protein interactions in the plant cell cycle has progressed in recent years, information on the SAC mechanism in this context remains scarce. Here, we used bigBac, chemical crosslinking, mass spectrometry and structural analysis to identify protein‒protein interactions (PPIs) among AtCDC20, AtBubR1 (BMF2) and AtMad2, which are components of the core MCC in *A. thaliana.* Fifty-three nonredundant crosslinks were identified, with multiple binding sites in different domains and subdomains of these proteins, such as the β-propeller (AtCDC20), tetratricopeptide domain, and KEN boxes, including a new KET box (AtBubR1) and Horma domain (AtMad2). We also identified a hierarchy of interactions between these three proteins, with the potential to regulate signalling through the SAC molecular mechanism. Interactions that characterize the Mad2 tetramer, with the coupling of two dimers, were also identified. No PPIs were identified for AtBub1 (BMF1) or AtBub3:1, revealing a significant difference in MCC formation compared with that in other eukaryotes. Our findings provide a structural basis for core MCC formation in *A. thaliana* and offer new insights into the proteins CDC20, BubR1 and Mad2 in the context of the SAC.

## Introduction

Almost seventy years ago, Alma Howard and Stephen Pelc first described the timeframe of cellular life according to four periods of cell division in root beans (Howard & Pelc, 1986), paving the way for understanding the eukaryotic cell cycle. However, this area of inquiry in plants is lagging behind the corresponding research in animals and yeast. Therefore, understanding the structural organization and dynamics of protein homo and heterocomplexes related to cell cycle control is important for overcoming this gap. Unlike what occurs in animals, environmental factors such as light intensity, temperature, water availability, and nutrient supply can cause significant changes in the cell cycle and development of plants (Stals, H., & Inzé, 2001). However, the basic molecular machinery controlling the cell cycle is largely conserved in eukaryotes (Kops et al., 2020; Dewitte & Murray, 2003; De Veylder et al., 2007). Understanding the molecular mechanisms that control the cell cycle during development could contribute to the development of resilient varieties for food production and thus mitigate the effects of ongoing climate change (Carneiro et al., 2021). The spindle assembly checkpoint (SAC) is a regulatory molecular mechanism executed mainly by the mitotic checkpoint complex (MCC) to monitor the anchoring of mitotic spindle microtubules to ensure correct chromosome segregation, and the core MCC is formed by three essential SAC components: CDC20, BubR1 and Mad2 (Musacchio 2015). Although our knowledge of protein‒protein interactions in the plant cell cycle has progressed in recent decades (Van Leene et al., 2011), information on the SAC mechanism in this context remains scarce. The cell division cycle 20 protein (CDC20) has a β-propeller domain formed by seven WD40 repeats (blades), each with four antiparallel β-strands (Kapanidou et al., 2017). Budding uninhibited by benzimidazole-related 1 (BubR1) has a tetratricopeptide (TPR) domain (D’Arcy et al., 2010) and conserved KEN boxes (Sczaniecka et al., 2008). Mitotic arrest deficient 2 (Mad2) is a HORMA protein (Rosenberg et al., 2015) with the capacity to assume two conformations, namely, open (O-Mad2) and closed (C-Mad2), after interacting with Mad1 and CDC20 (Luo et al., 2001; Sironi et al., 2001). Because many studies of SAC interactions involve peptides or truncated proteins, studies involving full-length proteins can provide new insight into this molecular mechanism because the PPI might have remained hidden under these experimental conditions. Chemical crosslinking and mass spectrometry (crosslinking-MS) enables the identification of protein‒protein interactions (PPIs) and heterocomplex characterization (O’Reilly and Rappsilber, 2018; Rozbeský et al., 2018; Graziadei et al., 2022; Botticelli et al., 2024). These techniques combined with baculovirus expression systems (Pan et al., 2018) and bigBac approaches (Weissmann et al., 2016) have become robust strategies for PPI identification. Thus, here, we combined these strategies to identify and structurally characterize the interactions between the proteins that form the MCC core in *Arabidopsis thaliana*, AtCDC20 AtBubR1 (BMF2) and AtMad2 (Fig. 1).

**Figure 1:**
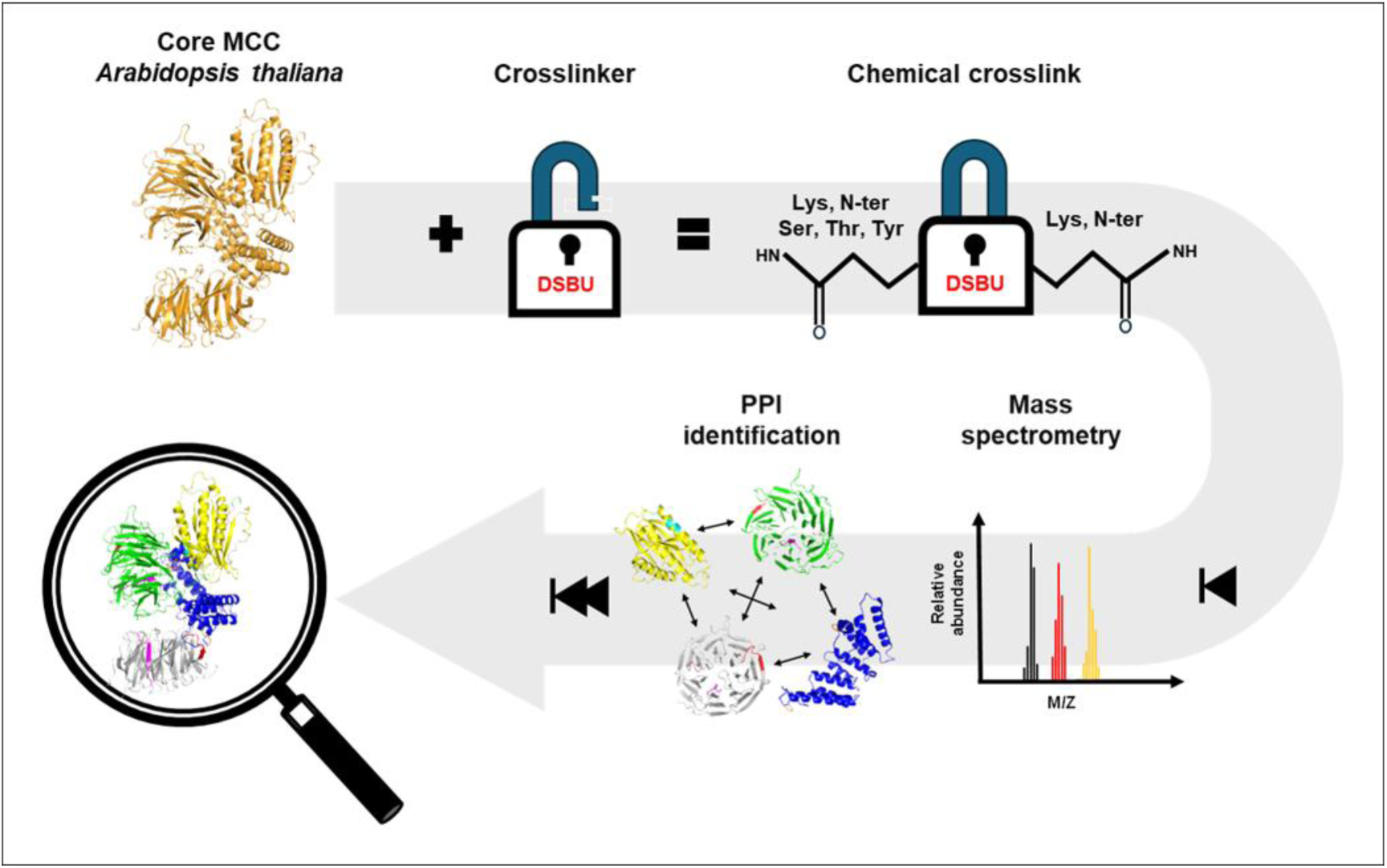
Graphical representation of the strategy for identifying protein‒ protein interactions (PPIs) involved in the formation of the MCC core in *A. thaliana*. First, the MCC core components (AtBubR1, AtCDC20, and AtMad2) are expressed and purified \**in vitro*\* as full-length proteins. Next, a cross-linking reaction is performed using disuccinimidyl dibutyric urea (DSBU). An enzymatic digestion protocol is subsequently applied, followed by peptide identification via mass spectrometry. The process concludes with the mapping of cross-links and a comparative analysis against structural models.

## Results

### *In vitro* formation and components of the core MCC of *Arabidopsis thaliana*

Insect cell culture combined with a baculovirus expression system allows us to produce high-quality cells (Trowitzsch et al., 2010). Therefore, using bigBac technology (Weissmann et al., 2016), six pLIB vectors with full-length proteins were constructed, pLIB MBP AtCDC20, pLIB 6his AtBubR1 (BMF2), pLIB 6his AtMad2, 6his AtBub1, 6his AtBub3;1 and pLIB 8TEV AtCDC20 (Fig. 2a), for which the pBIG1a MBP core MCC bigBac vector was employed (Fig. 2b). These constructs were then transfected into Sf9 cells to generate baculoviruses. Individually or as core MCCs in bigBac. An affinity column was used for the purification of the core MCC, and we observed three bands with expected sizes of approximately 92.2, 46.3 and 24.5 kDa for MBP AtCDC20, 6his AtBubR1 and AtMad2 6his, respectively (Supplementary Figure 1). Analogous results were obtained in the second purification step via size-exclusion chromatography (SEC) on a Superose 6 column, and the results were analysed by SDS‒PAGE (Fig. 2d–e). These results revealed that in *Arabidopsis thaliana,* the core MCC components (AtMad2, AtCDC20 and AtBubR1) interact reciprocally, as observed in yeast (Chao et al., 2012) and humans (Alfieri et al., 2016). In eukaryotes, the cell division cycle 20 protein (CDC20) is an essential component of the SAC. (Musacchio and Salmon, 2007, Tian et al., 2012). In *A. thaliana*, AtCDC20 has four hundred and fifty-seven amino acids, with a MIM motif in the N-terminal region and a short C-terminal region. Its central region is a classical β-propeller domain (Fig. 2e,f and Supplementary Figure 2), with seven WD40 repeats (blades) each formed by four antiparallel β-sheets stabilized by hydrophobic interactions (Tian et al., 2012; Xu & Mim, 2011; Yu, 2007). Our analysis of sequences from different plant species (Supplementary Figure 3) revealed a classic AtCDC20 MIM motif (MIM1) in the N-terminal region (R88-P96); two further MIM motifs were identified in the β-propeller domain, namely, a noncanonical MIM motif (MIM2) on blade 3 and, using the ELM tool (Kumar et al., 2020), we identified a third motif (MIM3) on blade 7 (R404-P412), as shown in Figure 2 e,f). In plants, compared with MIM2, MIM1 is more variable. However, MIM3 is conserved, suggesting its biological importance. Nevertheless, to function in a SAC context, the MIM3 motif must be accessible to Mad2. This requires transient and partial unfolding of the β-propeller domain. In this context, we identified a conserved sequence (G333-R339) referred to here as the glycine box (Fig. 2 e,f). Additionally, the structural characteristics of the β-sheets of blade 1 (E129-D133) and blade 7 (E424-W430) are shown in supplementary Figure 4. Both will be detailed later. In *Arabidopsis thaliana,* BubR1 (AtBubR1) has 396 residues and is composed of eight antiparallel α-helices, with the TPR domain in the N-terminal region (Fig. 2g, h) (D’Arcy et al., 2010; Bolanos-Garcia et al., 2011). This TPR domain presents six α-helices with a concave shape, forming subdomains TPR1 (P84-K105), TPR2 (L126-H152) and TPR3 (A159-R188). AtBubR1 has three classical KEN boxes: KEN1 (K31–N33) in the N-terminal region before the TPR domain and KEN2 (K220–N222) and KEN3 (K305–N307) after the TPR domain (Fig. 2g, h and Supplementary Figure 5 and 6). We also identified a new conserved KEN box (K376-T378) in which asparagine was replaced by threonine, named here the KET box. Only KEN box 2 is not conserved in plants. (Figures 2 g, h and Supplementary Figure 5). Mad2 is a HORMA protein (Rosenberg & Corbett, 2015; Aravind, L., & Koonin, 1998), and *Arabidopsis thaliana* (AtMad2) has 207 residues and a HORMA domain (S17–H192) (Figure 2 i,j and Supplementary Figure 7 and 8). We identified two regions in AtMad2 that mediate classical and no canonical interactions with AtCDC20, AtBubR1 and AtMad2, suggesting that it functions in SAC regulation. (Figure 2 i, j and Supplementary Figure 7 and 8). Both will be detailed later. Thus, our results reveal that even considering the difference in size of BubR1, the components of the core MCC of *A. thaliana* (AtMad2, AtBuBR1 and AtCDC20) exhibit the same general composition of domains and motifs for SAC functions, which has already been identified in other eukaryotes (Kops et al., 2020; Musacchio & Salmon, 2007; Hardwick et al., 2000). The complete MCCs characterized in other eukaryotes include the proteins Bub1 and Bub3 (Kops et al., 2020; Musacchio, 2015; Musacchio & Salmon, 2207; van Hooff et al., 2017). Therefore, to investigate whether this composition is also applicable to *A. thaliana*, we performed size-exclusion chromatography (SEC) on a Superose 6 column and performed pull-down experiments. In both experiments, we used the core MCC plus AtBub1 (BMF1) and AtBub3:1. In the case of the pull-down, amylose and MBP-AtCDC20 (as bait) were used. The two experiments were analysed by SDS‒PAGE. Our combined pulldown and SEC results revealed that neither AtBub1 nor AtBub3;1 bind the core MCC (Fig 3 a, b and Supplementary Figure 9). These results corroborate those of previous studies in *A. thaliana* (Komaki and Schnittger, 2017). Chemical crosslinking (Pan et al., 2019), when combined with bigBac approaches, becomes an efficient strategy for *in vitro* studies of protein complex formation. Therefore, to identify protein‒protein interactions (PPIs) between AtCDC20, AtMad2, and AtBubR1 in the core MCC of *A. thaliana*, we performed chemical crosslinking (Fig. 3c–h) with disuccinimidyl dibutyric urea (DSBU) (Pan et al., 2018). Different combinations of full-length proteins were used: a) core MCC with MBP AtCDC20 plus AtBub3:1 and AtBub1; b) just core MCC with MBP AtCDC20; c) MBP AtCDC20 plus AtMad2; and d) core MCC with 8TEV AtCDC20 (Fig. 3 c-h). A total of six thousand and five hundred and twenty-four interactions were identified, of which sixty-three were nonredundant crosslinks. Among these three proteins, twenty-five are intramolecular, and thirty-eight are intermolecular. Nonredundant crosslinks were used to validate the models (Chao et al., 2012; Alfieri et al., 2016; Mapelli et al., 2007) applied here (Supplementary Tables 1–6). The nonredundant cross-links C3–C7, C12–C17, C32–C36, C40, C41, B1-B7, and M1-M7 confirm the composition of the MCC core in *A. thaliana* with AtMad2, AtCDC20 and AtBubR1 and reciprocal interactions between these proteins (Fig. 3a–n; Supplementary Fig. 10-13 and Supplementary Tables 1–6), corroborating the results of studies in other eukaryotes. Chao et al., 2012; Alfieri et al., 2016; Mapelli et al., 2007). We identified the hierarchical order of the interactions between Mad2 and the other genes (Fig. 3 I, k and n). In the presence of AtBubR1, residues K114 and K190 of AtMad2 interact with residues K376 and T378 on the KETbox of AtBubR1 (Fig. 3i,n and supplementary table 5). However, in the absence of AtBubR1, residues K114, S117, and K183 of AtMad2 interact with residue K221 of AtCDC20 (Fig. 3k and Supplementary Table 4). This interaction site corresponds to MIM2 e and is shown in the nonredundant cross-links C41–C43 and B9–10 (Fig. 3j, Supplementary Fig. 7 and Supplementary Table 4). The AtMad2 sequence identified here as a KIM (for KET box Interact Motif) with a preferred affinity for binding to BubR1 (I105–R118) includes a conserved sequence (E110–R118) corresponding to T109–R117 in human cells. To create “stable” O-Mad2 (Mad2^LL^), this sequence was deleted (Mapelli et al., 2007). Thus, our findings suggest that AtBubR1 can block this region and thereby regulate the stoichiometry of O-Mad2 and C-Mad2 in a SAC context. Future studies will be necessary to confirm or refute this hypothesis. Additionally, the alignments show that these cross-linked regions involved in this hierarchy of interactions are conserved in plants, corroborating our hypothesis (Fig. 3 I, k, n and Supplementary Figures 3, 5 and 7).

**Figure 2:**
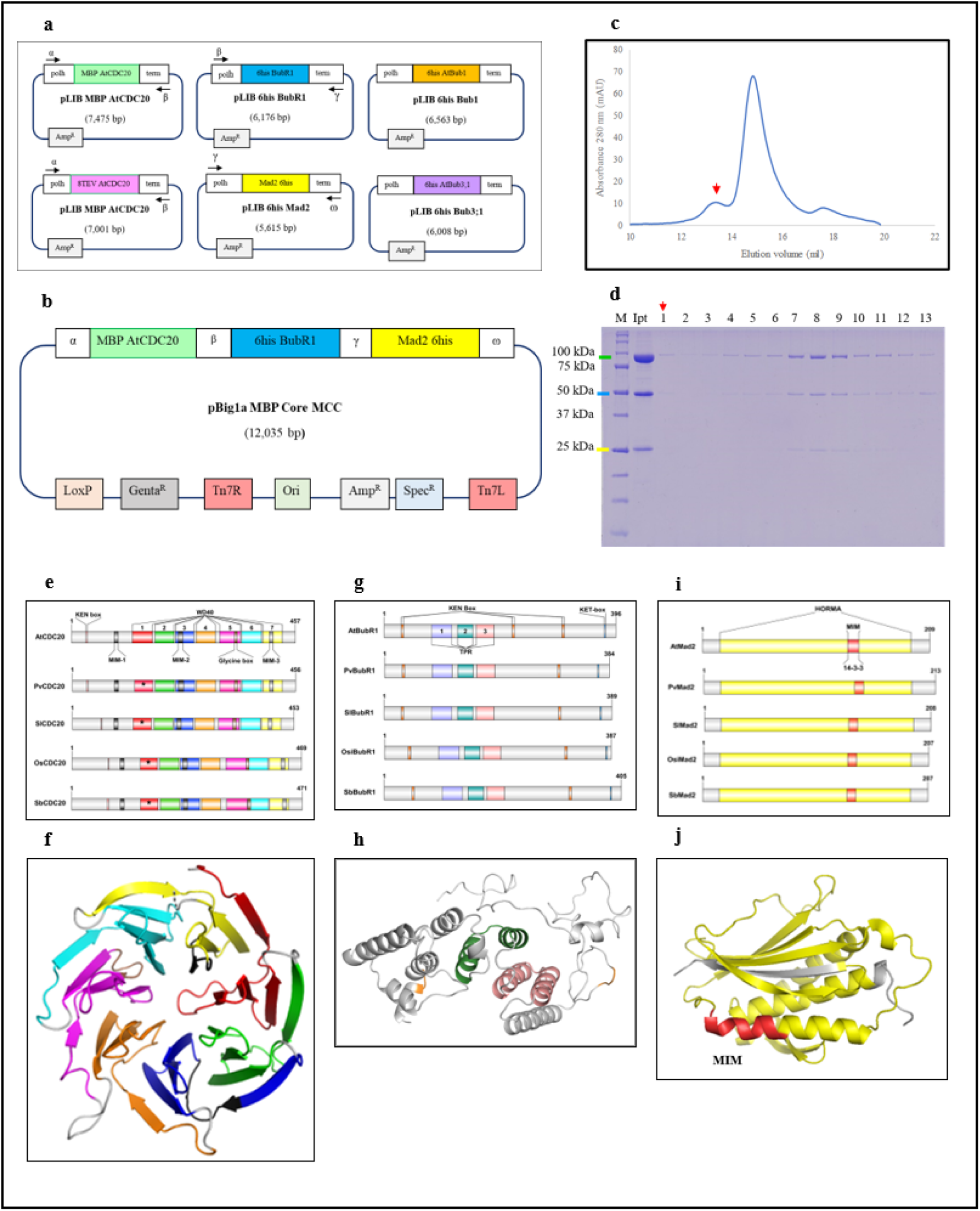
Expression, purification and structural analysis of core MCC proteins of *Arabidopsis thaliana*. **a,** Schematic representation of the pLIB vector constructs of SAC proteins, including MBP AtCDC20 (green), orange (6his-AtBub1), 6his AtBubR1 (blue), AtMad2 6his (yellow), AtBub1 6his (pink) and AtBub3:1 6his (lilac). **b**, Schematic representation of the bigBac assembly of the *Arabidopsis thaliana* core MCC proteins, including MBP AtCDC20 (green), 6his AtBubR1 (blue) and AtMad2 6his (yellow). **c**, Size-exclusion chromatography with core MCC proteins (MBP AtCDC20, 6His AtBubR1 and 6His AtMad2) using Sepharose S6 and isocratic conditions. Red arrows indicate only AtCDC20 and AtBubR1 in a complex. **d**, SDS‒PAGE analysis of size-exclusion chromatography with sample A (core MCC proteins). Red arrows indicate only AtCDC20 and AtBubR1 in a complex. The coloured bars indicate the relative positions of the SAC proteins: green (MBP-AtCDC20), blue (6-His-AtBubR1), and yellow (6His-AtMad2). The red arrow indicates a slot with only the MBP AtCDC20 and 6his AtBubR1. **e**, Cartoon representation of the conserved topology of plant CDC20 showing the KEN box (salmon), MIM motifs (black), seven WD40 repeat blades: 1 (red), 2 (green), 3 (blue), 4 (orange), 5 (magenta), 6 (cyan) and 7 (yellow) and the glycine box (brown). **f**, Cartoon representation of the β-propeller of AtCDc20 showing seven WD40 repeat blades, MIM motifs, and glycine boxes. The colour scheme is the same as that in Figure 2e. N- and C-termini were omitted. **g**, Cartoon representation of the AtBubR1 N-terminal region topology with KEN box 1 and KEN box 2 (orange) separated by the conserved TPR domain composed of TPR1 (lilac), TPR2 (green), and TPR3 (salmon). **h**, Cartoon representation of the TPR domain of AtBubR1. The colour scheme is the same as that in Figure 2g. **i**, Cartoon representation of the conserved topology of AtMad2 with the HORMA domain (yellow) and the MIM motif (red). **j**, Cartoon representation of AtMad2: HORMA domain. The colour scheme is the same as that in Figure 2i. Plant species: At (*Arabidopsis thaliana*), Pv (*Phaseolus vulgaris*), Sl (*Solanum lycopersicum*), Os (*Oryza sativa*) and Sb (*Sorghum bicolor*).

**Figure 3:**
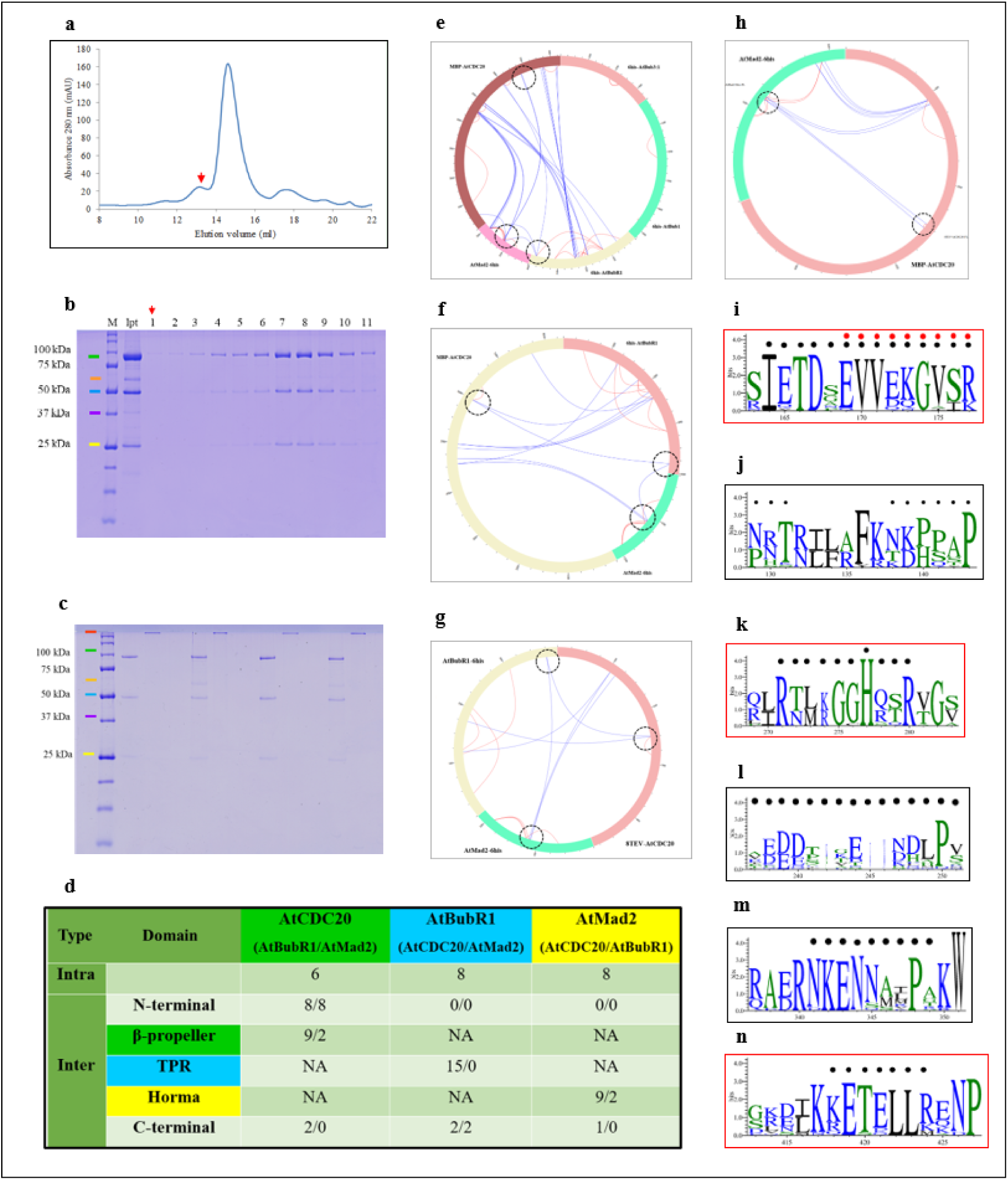
Protein‒protein interactions for core MCC formation in *Arabidopsis thaliana* in *Arabidopsis thaliana*. **a,** Size-exclusion chromatography with core MCC proteins plus AtBubu3:1 and AtBub1. **b**, SDS‒ PAGE analysis of size-exclusion chromatography with the core MCC proteins plus AtBubu3:1 and AtBub1. The coloured bars indicate the relative positions of the SAC proteins: green (MBP-AtCDC20), orange (6his-AtBub1), blue (6-his-AtBubR1), purple (6his-Bub3:1), and yellow (6his-AtMad2). **c**, SDS‒PAGE analysis of DSBU crosslinking results with SAC proteins. 1 without and 2 with DSBU. The coloured bars indicate the relative positions of the SAC proteins: red (cross-linked proteins), green (MBP-AtCDC20), orange (6his-AtBub1), blue (6-his-AtBubR1), purple (6his-Bub3:1), and yellow (6his-AtMad2). **d**, Table 1, Summary of the nonredundant crosslinks obtained in experiments A, B, C and D, distributed according to different regions of the proteins. **e**, Circular plot showing the nonredundant crosslinks obtained in experiments with MBP-AtCDC20, 6His AtBubR1, AtMad2, AtBub1 and 6His AtBub3:1 (A); **f** Circular plot showing the nonredundant crosslinks obtained in experiments with MBP-AtCDC20, 6His AtBubR1 and AtMad2 (B). **g**, Circular plot showing the nonredundant crosslinks obtained in experiments with 8TEV-AtCDC20 and AtMad2-6his (C). **h**, Circular plot showing the nonredundant crosslinks obtained in experiments with 8TEV-AtCDC20,6-HHis- and AtMad2-6His (D). Red and blue lines indicate intra- and intermolecular nonredundant crosslinks, respectively. The dashed black circle in Figure 3 e-h indicates the regions of AtCDC20, AtMad2 and AtBubR1 that participate in hierarchical interactions. Consensus sequence logos with interaction motifs included **i** KIM (AtMad2), **j** MIM1 (AtCDC20), **k** MIM2 (AtCDC20), **l** KEN box 2 (AtBubR1), **m** KEN box 3 (AtBubR1), and **n** KET box (AtBubR1). The black circles indicate the amino acids included in the peptide identified in the crosslinking experiment. Red circles indicate a conserved sequence that, in HsMad2, has been replaced by a GSG to generate stabilized O-Mad2 (Mapelli et al., 2007). Red boxes indicate regions containing peptides identified within the hierarchical interaction between AtMad2, AtBubR1 and AtCDC20.

### Homocomplexes of AtMad2 in *Arabidopsis thaliana*

Mad2 plays a central role in SAC signalling because of its ability to switch its structure from open to closed (Luo et al., 2000; Sironi et al., 2001) to form homocomplexes as dimers (Mapelli et al., 2007) or tetramers (Fang et al., 1998). Our lack of redundant crosslinks B9, B10 and C9-43 and C28–C32 revealed that in *A. thaliana*, Mad2 can interact with both CDC202 and BubR1 in its central and C-terminal regions (Supplementary figure 10-14 and Supplementary tables 4 and 5). In the dimer of AtMad2, the sequence delimited by R134-Y142 has a substantial hydrophobic content (Supplementary Figure 7). The residues R134, S138, T141 and Y142 of C-Mad2 can form strong hydrogen bonds (≤ 3 Å) with Y34, N30, P53 and Q130 of I-Mad2 (Fig. 4a–d), analogous to hMad2 R133, Q134, T136 and T140 (Hara et al., 2015; Mariani et al., 2012). In human cells, T136, A137, T140 and F141 (S137, S138, T141 and Y142 in *A. thaliana*), located at the asymmetric contact interface of hMad2, have implications for conformational change (Mapelli et al., 2007; Hara et al., 2015; Mariani et al., 2012). The sequence of residues that interact with Mad2 and has implications for the functionality of the SAC, is called an MIM motif (Hara et al., 2015; Mariani et al., 2012). Therefore, by definition, the interaction region in the AtMad2 dimer is composed of the amino acids R134–Y142 (R133–F141 in hMad2) and is a noncanonical MIM motif (Fig. 4c,d). Additionally, the observation of crosslinks with DSBU and measured distances between K114, S117, and K183 (AtMad2) and K221 (AtCDC20) above forty angstroms suggests that these interactions are possible only with an AtMad2 dimer (or tetramer) in the core MCC. The implications of the core MCC composition involving the dimer will be discussed later. Although the structure and formation of Mad2 dimers have been studied (Mapelli et al., 2007; Hara et al., 2015; Mariani et al., 2012), aspects of tetramer formation remain unknown. In human cells, tetramers of Mad2 inhibit cell cycle progression *in vitro* through the inhibition of APC/C activity (Fang, et al., 1998). Crosslinks M8 among amino acids S61 and K123 and M9 and M10 (K92–K114 and K92–S117, respectively) revealed that the distance between the residues “should” be greater than 12.5 Å (Fig. 4g, Supplementary Figure 14). We used O-Mad2 in model 2v64, where K114 and S117 are missing. (Mapelli et al., 2007). N109 and E110 are shown as a reference for a hypothetical position of these amino acids (Supplementary Figure 14). In this case, our data revealed that a tetramer of AtMad2 formed with two dimers docked in an overlapping arrangement. The MIM motif is a contact point between AtMad2 dimers (Fig. 4a–c) and is composed mostly of hydrophobic residues (Fig. 4a–c) and Supplementary Figure 10). Thus, the AtMad2 tetramer forms a primary hydrophobic core delimited longitudinally by four arginine (R134) residues (Fig. 4a–h).

**Figure 4:**
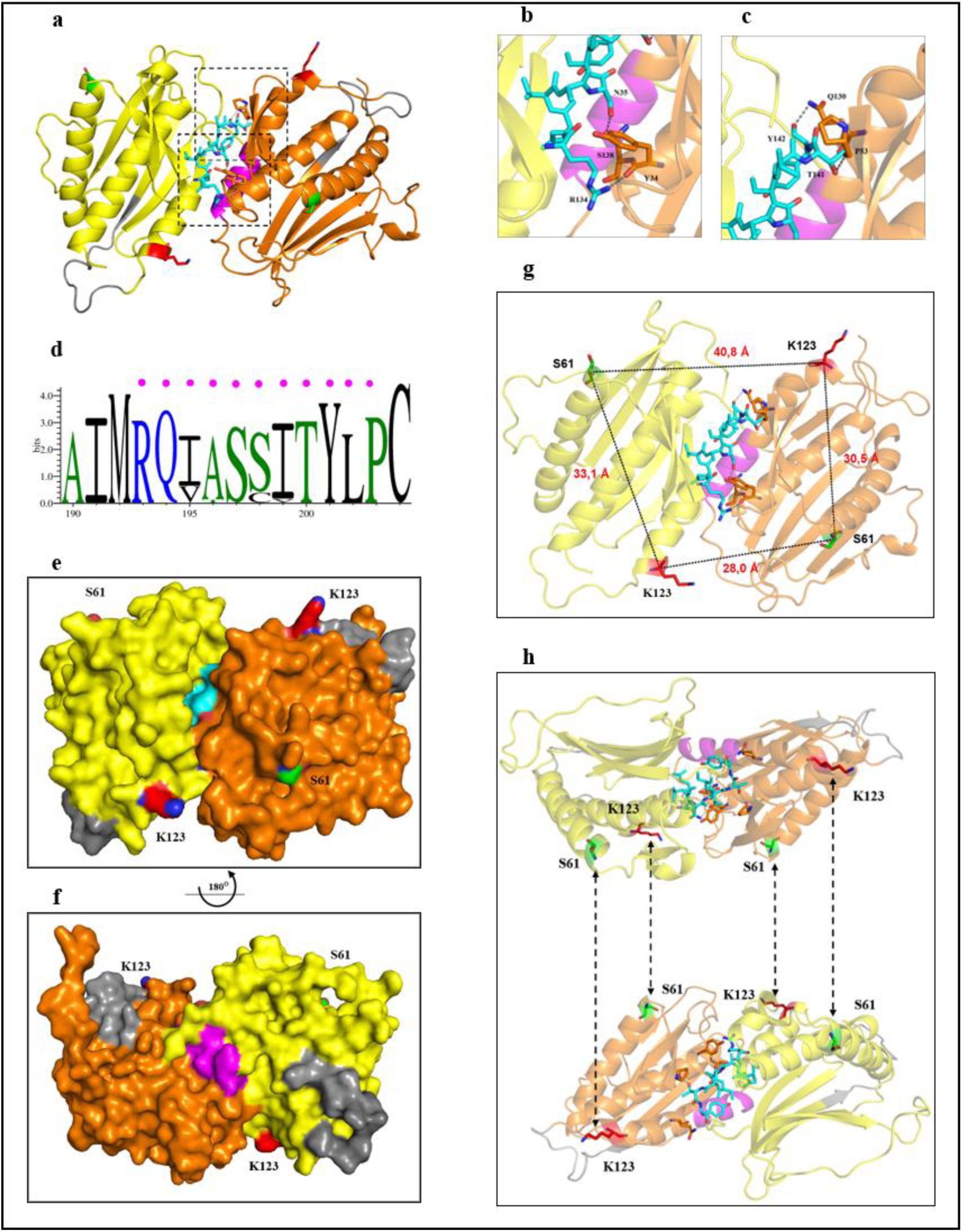
Mad2 dimer and tetramer structure in *Arabidopsis thaliana* based on crooslink experiments. **a,** Cartoon representation showing an overview of the Mad2 dimer. O-Mad2 (orange and MIM motif magenta), C-Mad2 (yellow and MIM motif cyan) and “safety belt” (black). **b and c**, Cartoon representation showing detailed views of the interaction motif (MIM motif) in the AtMad2 dimer interface. The colour scheme is the same as that in Figure 4a. Colours elements: nitrogen (blue), oxygen (red) and potential hydrogen bond formation (≤ 3 Å) between the amino acids of the primary hydrophobic core are dashed black lines; **d**, Consensus sequence logos with AtMad2’s MIM motifs included; pink circles indicate a prediction for 14-3-3 phosphorylation; **e** and **f**, Surface view from the bottom and top, respectively, with 180° rotation and exposed amino acids S61 and K123. Colours elements at S61 and K123: nitrogen (blue) and oxygen (red); **g**, Cartoon representation of the Mad2 dimer showing the distances in angstroms between amino acids S61 and K123; **h**, Cartoon representation of the proposed model of AtMad2 tetramer formation with the interaction point between the two Mad2 dimers on S61 and K123, indicated by black dashed lines. The model used 2v64 (Mapelli et al., 2007).

### Structure of the core MCC

As observed in yeast and humans (Chao et al., 2012; Alfieri et al., 2016) and shown previously here. In *A. thaliana*, the core MCC is also composed of the proteins Mad2, BubR1, and CDC20. However, in plants, the interactions between the components of the core MCC have been much less studied than those in other eukaryotic models and, consequently, are poorly understood. Two Cdc20 binding sites on BubR1 were demonstrated in studies with human cells, with implications for SAG regulation. (Davenport et al., 2006). The lack of redundant C9-C25 crosslinks (Supplementary Table 4) indicates the formation of a complex of AtBubR1 and two AtCDC20 molecules (Fig. 5a–i and Supplementary Table 4), referred to here as CDC20 of mitotic complex 2 (CDC20^MC1^ and CDC20MC2). Interestingly, the concentration of AtMad2 does not appear to be sufficient to interact with all AtBubR1 and AtCDC20 molecules. In this context, the nonredundant crosslinks (C9-C11, C18, C19, C26-C37, C39, B10, B11, and M8-M10) revealed that some interaction sites between CDC20 and BubR1 are also shared by AtMad2 (Supplementary Tables 4 and 5). Thus, the absence of AtMad2 enables the interaction of multiple AtCDC20 and AtBubR1 units. This explanation is corroborated by the observation of SEC/S6 in slot 1, corresponding to AtCDC20 and AtBubR1 complexes with a molecular mass greater than the core MCC in slots 3-13 (Fig. 1c, d and Fig. 2a, b). We speculate that the presence of AtMad2 predominantly in the form of dimers or tetramers explains this finding. Further investigations will be needed to determine the stoichiometry of the interactions between AtCDC20 and AtBubR1 in the presence and absence of AtMad2 homocomplexes. In human cells, APC/C and Mad2 compete for KILR sites in CDC20 (Izawa & Pines, 2012; Overlack et al., 2015). The presence of nonredundant crosslinks (C13–C16 and C26–C36) revealed that AtMad2 and AtBubR1 bind AtCDC20 similarly in the conserved motifs R88, I89, L90 and A91 (Supplementary Tables 4 and 5). These findings suggest that AtBubR1 regulates the interaction between AtMad2 and AtCDC20 at this site. AtCDC20 binds the C-terminus of AtBubR1 or AtMad2 at the top of the β-propeller and shows conserved interactions (Izawa & Pines, 2012; Overlack et al., 2015). In contrast to KEN box 2, KEN boxes 1 and 3 are conserved in plants (Fig. 2g; Fig. 3m, n Fig. 6f and Supplementary Fig. 5). Our results revealed that AtCDC20 bound AtBubR1 in the N-terminal and C-terminal regions, as well as the TPR3 domain (Supplementary Figures 10, 11 and 13 and Supplementary Tables 1 and 2). At the C-terminus, AtCDC20 interacts with AtBubR1 on KEN box 3 and the KET box (Supplementary Figure 10 and Supplementary Tables 1 and 2). AtCDC20 also bound AtMad2 in the Horma domain in the central and C-terminal regions (Supplementary Figure 10, Supplementary Table 3). The crosslinking of C13 (Supplementary Table 4) suggests the *in vitro* conditions under which AtBubR1 itself can block the RILA-mediated impairment of binding between AtMad2 and AtCDC20 (C9–C11 and C26–C29). This involves hotspot interaction sites of KEN boxes (C23–C25 and B1–B8) with possible implications for APC/C activity (King et al., 2007; Keskin et al., 2005). and autoinhibitory binding with a second CDC20 in the functional SAC (Tipton et al., 2001). Examples are provided by the nonredundant crosslinking of B1 between the first and second KEN boxes or B7 between the second and third KEN boxes with the new KET box (Fig. 3e–g, l-n; Supplementary Figure 12; Supplementary Table 5). As previously mentioned, our ELM analyses revealed an MIM motif (MIM3) in AtCDC20 on blade 7 (Fig. 2 e, f). However, MIM3 is buried within the β-propeller, preventing access by AtMad2 (Supplementary Figure 4). It has been demonstrated that the absence of the “velcro” formed by hydrophobic interactions between the β-sheets of blades 1 and 7 can allow for a transient structural state, granting substrates access to the central cavity of the β-propeller domain. (Xu and Mim., 2011). This feature is found in AtCDC20, where these hydrophobic interactions between blades 1 and 7 are unlikely (Fig. 6a, b). Additionally, we identified a conserved glycine box (G333–R339) in AtCDC20 that may facilitate partial unfolding of the β-propeller domain because of its intrinsic flexibility (Fig. 6c–e). Studies have already shown unfolding and refolding of the β-propeller for Tc toxin activation (Gatsogiannis et al., 2018). In a SAC context (Chao et al., 2012), this glycine box can form hydrogen bonds with both AtBubR1 in KEN box 1 (E32) and the AtMad2 dimer interface (R134, S138 and T141). However, in the interaction of the core MCC with the APC/C (Alfieri et al., 2016), these hydrogen bonds are unlikely (Fig. 6c–f). As previously shown, the Mad2 dimer interface is an MIM motif (Fig. 4a–d). Considering the measured distances and the models applied here, interactions between K221 (AtCDC20) and K114, S117 and K183 (AtMad2) of the nonredundant crosslinks C38, C42 and C43 are unlikely (Fig 7a, b and. They would only be possible through the coupling of a dimer (or Mad2 tetramer). This would place at least two MIM motifs (antiparallel) of AtMad2 near the glycine box (AtCDC20) and the KEN box of AtBubR1 (Fig. 6 a, b and Supplementary Table 4). Thus, all the difficulties involved in representing a three-dimensional structure in a two-dimensional illustration should be considered. We hypothesize a model for the composition of the core MCC in *Arabidopsis thaliana*, featuring AtMad2 as a homocomplex (dimer or tetramer) and two AtCDC20 molecules and using AtBubR1 as a platform for both proteins (Figure 6 c). Its composition not only reflects features unique to this plant species but also provides new perspectives regarding core MCC organization in other organisms.

**Figure 5:**
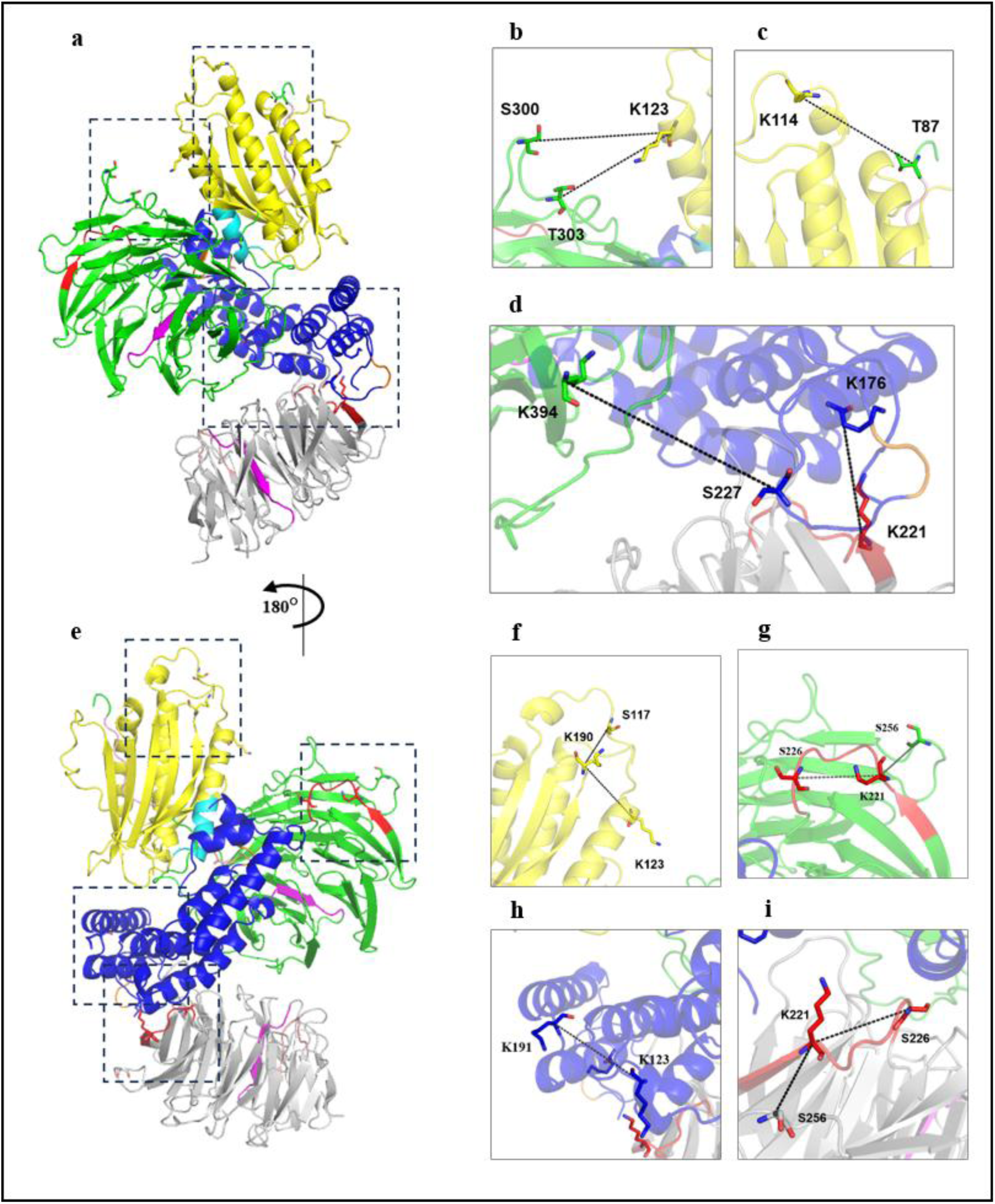
Positioning of two AtCDC20 molecules that interact with AtBubR1 and AtMad2 for core MCC formation in *Arabidopsis thaliana*. **a,** Cartoon representation of two AtCDC20 molecules, AtBubR1, and AtMad2, based on cross-linking experiments. AtCDC20^MC1^ (green), AtCDC20^MC2^ (gray), AtMad2 (yellow) and BubR1 (blue). MIM2 (red), MIM3 (magenta), glycine box (salmon), AtMad2’s MIM (cyan), and KEN boxes 1 and 2 of AtBubR1 (orange) of AtCDC20. **b,** Intermolecular crosslinking between AtCDC20^MC1^ (S300 and T303) and AtMad2 (K123); distance measured in angstroms, 18,7 Å and 16,9 Å, respectively; **c,** intermolecular crosslinking between AtCDC20^MC1^ (T87) and AtMad2 (K114), 17,2 Å; **d,** intermolecular crosslinking between AtCDC20^MC1^ (K394) and AtBubR1 (K227) and between AtCDC20^MC2^ (K221) and AtBubR1 (K176), 22,4 Å and 14,2 Å, respectively. **e,** Cartoon representation of two AtCDC20 molecules, AtBubR1, and AtMad2, based on crosslinking experiments with 180° rotation and using the same colour scheme as in Figure 5a. **f,** Intramolecular crosslinking between amino acids S123, K123 and K190 (AtMad2), 9,1 Å and 13,1 Å, respectively. **g,** Intramolecular crosslinking between amino acids K221, S226 and S256 (AtCDC20 ^MC1^), 10,6 Å and 10,3 Å, respectively. **h,** Intramolecular crosslinking between amino acids K123 and K191 (AtBubR1), 14,1 Å. **i,** Intramolecular crosslinking between amino acids K221, S226 and S256 (AtCDC20 ^MC2^), 10,5 Å and 10,0 Å, respectively. Colours: nitrogen (blue), oxygen (red). The 5LCW model was used (Alfieri et al., 2017).

**Figure 6:**
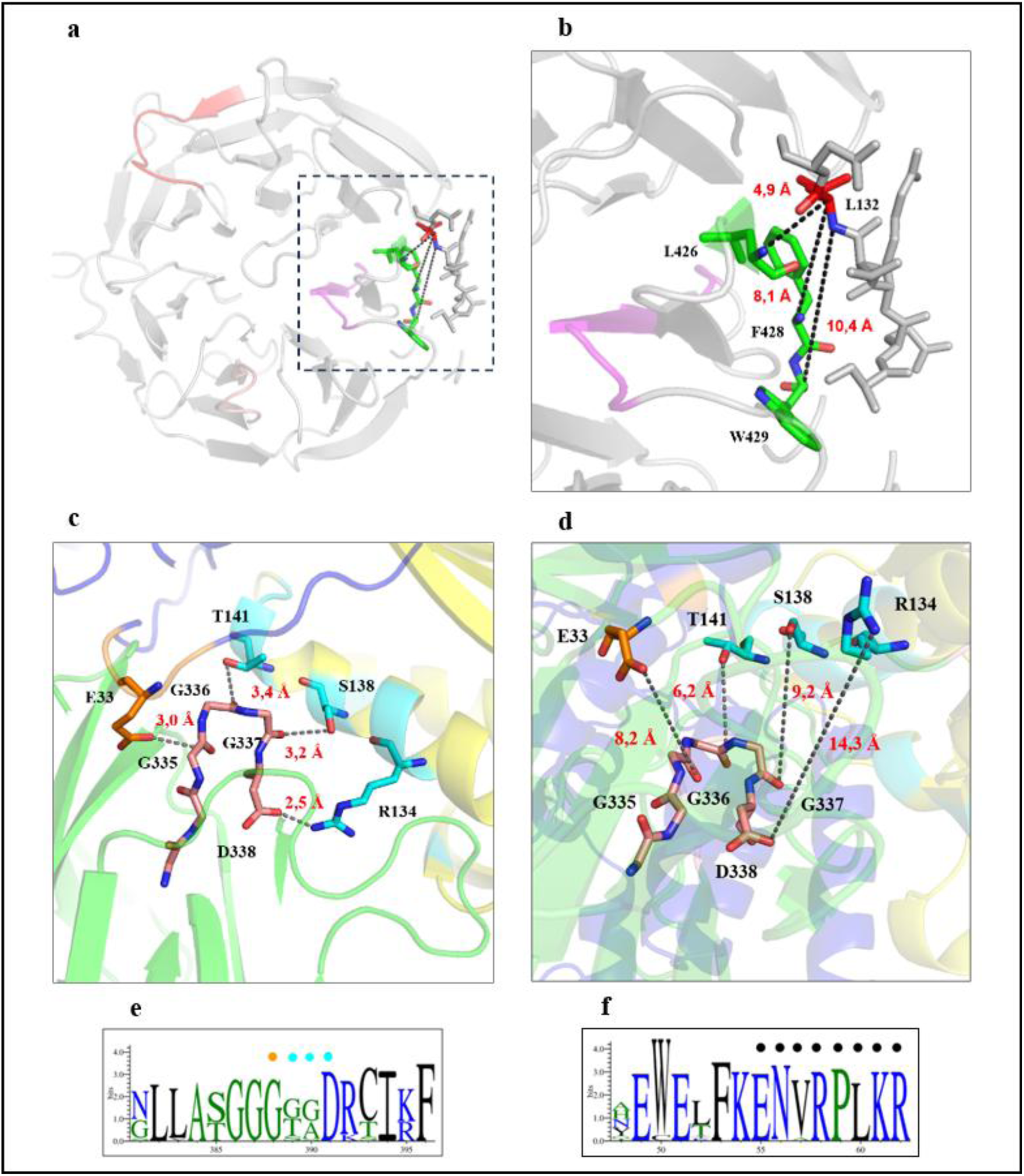
Characteristics of the AtCDC20 protein that enable structural flexibility of the β-propeller domain. **a**, Cartoon representation of the β-propeller of AtCDc20 showing seven WD40 repeat blades (gray); AtCDC20’s MIM2 and MIM3; red and magenta, respectively; the amino acid L132 (β-sheet of blade1) is shown in red; and the amino acids L426, F428 and W429 (β-sheet of blade 7) are shown in green. Amino acid shown in stick **b**, Detailed views of the amino acid L132 (β-sheet of blade1) shown in red; the amino acids L426, F428 and W429 (β-sheet of blade 7) are shown in green. Colours elements: nitrogen (blue), oxygen (red), the distances in angstroms between hydrophobic amino acids (red): L132 and the amino acids L426, F428 and W429 are dashed black lines. **c,** Detailed views of the amino acids with potential hydrogen bond formation between components of the core MCC in the context of the SAC: E33 KEN box1 (AtBubR1 in orange); G335, G336, G337 and D338 glycine boxes (AtCDC20 in salmon); and R134, S138 and T141 AtMad2’s MIM (cyan). The distances in angstroms are shown in red. The model used 4aez (Chao et al., 2007). **d**, Detailed views of the amino acids with unlikely hydrogen bond formation between components of the core MCC in the context of APC/C: E33 KEN box1 on AtBubR1 (orange); G335, G336, G337 and D338 glycine boxes (salmon); and R134, S138 and T141 AtMad2’s MIM (cyan). The distances in angstroms are shown in red. The 5LCW model was used (Alfieri et al., 2017). **e**, Consensus sequence logos with the glycine box included; amino acids with potential hydrogen bonds are indicated by coloured circles, with E33 on KEN box 1 at AtBubR1 (orange) or with R134, 138 and T141 on the AtMad2 of the MIM (cyan). **f**, Consensus sequence logos with KEN box 1 (AtBubR1). The black circles indicate the amino acids included in the peptide identified in the crosslinking experiment.

**Figure 7:**
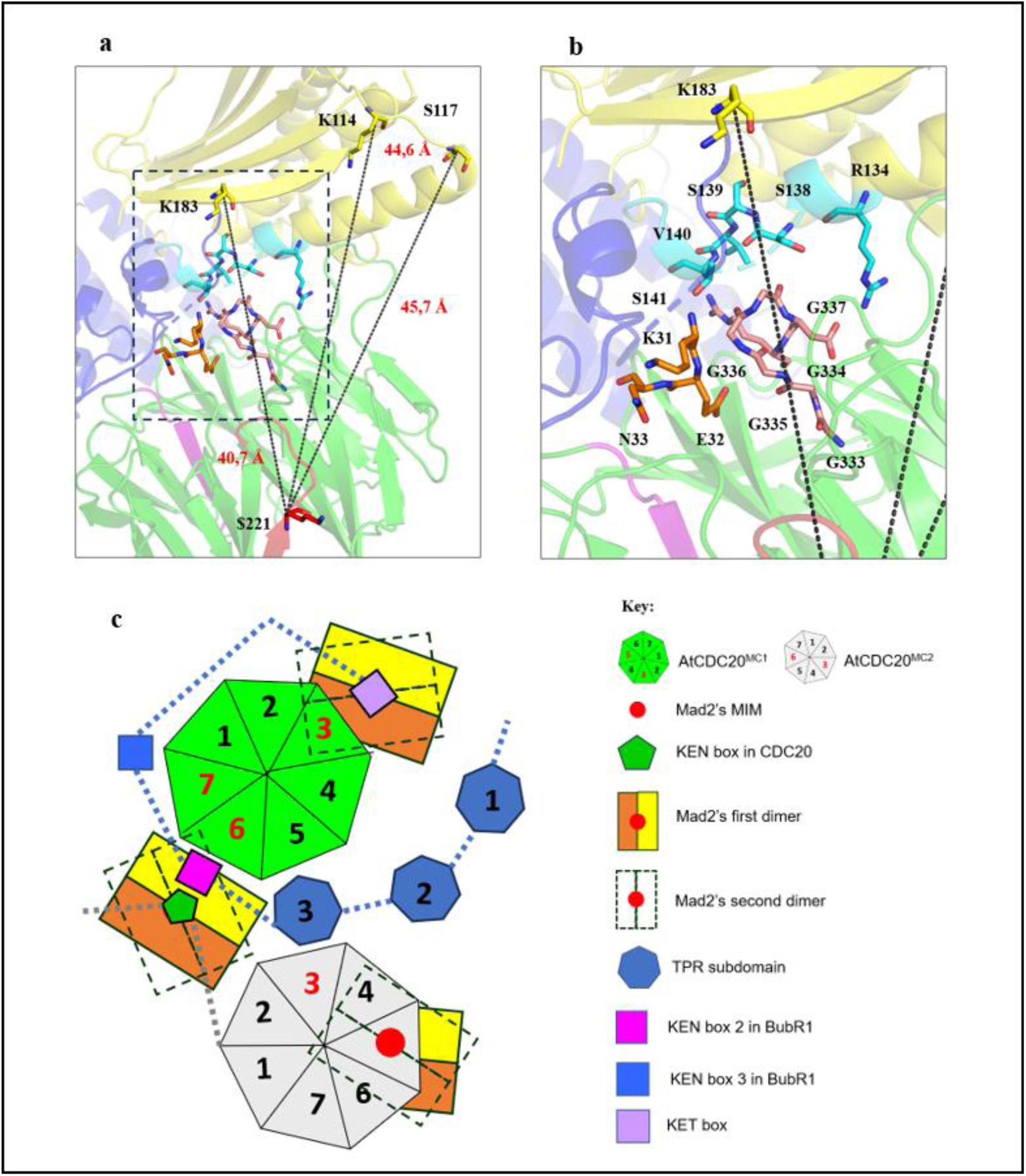
Positioning of the AtMad2 dimer in the core MCC of *Arabidopsis thaliana*. **a**, Cartoon representation showing measurements of the distances between amino acid S221 of AtCDC20 (red) and amino acids K114, S117 and S183 of AtMad2 (yellow). The measurements in angstroms (dashed black lines) are shown in red. Amino acids are also shown: KEN box 1 of AtBubR1 (orange), AtMad2’s MIM (cyan) and the glycine box of AtCDC20 (salmon). **b**, Detailed views showing the amino acids K31, E32 and N33 (KEN box 1 of AtBubR1) in orange; R134, S138, S139, V140 and S141 of AtMad2’s MIM (cyan); and G333, G334, G335, G336 and G336 (Glycine box of AtCDC20) in salmon. **c**, Schematic model for the composition and organization of the core MCC in *Arabidopsis thaliana* on the basis of the crosslinking results.

## Discussion

Plants modulate the development of their body plan by mediating signals from the immediate environment. These signals can be positive, such as nutrient availability (Liu and Lee., 2022), or originate from biotic and abiotic stresses (Zhu, J. K., 2016; Huang et al., 2019). This process requires the convergence of different signalling pathways linked to various phases of the cell cycle (Shimotohno et al., 2021) to synchronize the duplication and segregation of genetic material with conditions favourable for development. Despite significant advances in recent decades (Sablowski, R., & Gutierrez, C., 2022), our understanding of plant cell cycle control lags behind what is known about other eukaryotes. Adopting an evolutionary perspective broadens the horizons of our research. Identifying convergent and divergent aspects makes it possible to trace details that are otherwise difficult to observe in specific study models. This enables us to overcome barriers in understanding complex molecular mechanisms and points towards alternative methodological approaches. In this context, identifying intermolecular interactions and understanding the formation of protein complexes involved in the control of chromosome segregation in plants is important for elucidating the functions of these interactions, whether they are conserved or not (Liu et al., 2022). Our results revealed that in *A. thaliana*, the CDC20 and Mad2 proteins, with respect to size, domain composition and cell cycle-related interaction motifs (Fig. 1e–j; Supplementary Figures 3-9), are conserved (McAinsh and Kops, 2023). The cross-links of the N-terminal region of AtCDC20 with those of both AtMad2 and AtBubR1 are consistent with previous findings in related studies with canonical interactions, such as those involving the MIM motif, β-propeller domain, KEN-boxes and the TPR domain (Sironi et al., 2001; Tang et al.; 2001; King et al., 2007; Lara-Gonzalez et al. 2011 35–38). For example, studies have shown that CDC20 can interact with Mad2 on R88 and E170 (Chao et al., 2012) and crosslink C28-C32 on a “safety belt”, indicating that AtCDC20 can interact with AtMad2 in the same form (Fig. 4d–n and Supplementary Figures 10-14). Compared with HsBubR1 (McAinsh and Kops, 2023), AtBubR1 presents significant differences in length and domain composition (Fig. 2 g, h; Supplementary Fig. 10). In human cells, BubR1 is phosphorylated at different sites (on the C-terminal domain) related to microtubule attachment and chromosome congregation (Huang et al., 2008; Elowe et al., 2009). AtBubR1 (BMF2) is minor and does not have these sites, suggesting that it does not play a role in the docking of microtubules to the kinetochore, corroborating the findings of Komaki et al. (2017). Our findings suggest that in *A. thaliana*, AtBubR1 functions only in MCC assembly and SAC signalling (Elowe et al., 2009). Although the N-terminal region of AtBubR1 presents the same structural characteristics observed in human cells and yeast (D’Arcy et al., 2010 and Bolanos-Garcia et la., 2011), how the TPR domain and KEN boxes (Fig. 2g,h) are present remains unclear. In eukaryotes, complete and functional MCCs include Bub1 and Bub3 (McAinsh & Kops, 2023). Our experiments revealed that in *A. thaliana,* Bub1 (BMF1) and Bub3:1 do not bind to the protein complex formed by AtCDC20, AtMad2 and AtBubR1 (Fig. 1c). The GLEBS motif is related to the interaction with Bub3 in the WD40 of the beta propeller domain not found in AtBubR1 (Fig. 2 g, h). Therefore, even though it is relatively small, AtBubR1 (BMF2) is sufficient for core MCC formation and does not act to recruit AtBub1 and AtBub3.1 (Figs. 3 and 5). Compared with yeast and animal SAC signalling, MCC assembly is significantly different (McAinsh and Kops, 2023). However, further studies are needed to identify whether crucial underlying factors can (even if transiently) promote the association of the proteins Bub1 and Bub3:1 with the MCC core in *A. thaliana*. Studies on MCC formation in human cells have shown that the Mad1-Mad2 and Mad1-Bub1 complexes are required as physical components of the catalytic mechanism of formation (Faesen et al., 2017). Our results show that Mad1 and Bub1 (BMF3) are not necessary for *in vitro* core MCC formation in *A. thaliana*. Thus, the role of these proteins in MCC functionality in plants requires investigation. An essential and pivotal characteristic of Mad2 is its ability to undergo a topological switch, in which two different conformations, namely, open (O-Mad2) and closed (C-Mad2), after interaction with Mad1 and CDC20 on the MIM motif (Sironi et al., 2001; Kim et al. 2002; Yu et al., 2025 Luo et al., 2002). Another important characteristic of Mad2 is the formation of homocomplexes: dimers (Mapelli et al., 2006; Fang et al., 1998) and tetramers (Fang et al., 1998). Disruptions in Mad2 dimer and tetramer formation affect SAC progression in eukaryotic cells. (Mapelli et al., 2006; Fang., 1998; Mapelli et al., 2006) bition of APC/C activity. Therefore, we aimed to analyse protein‒protein interactions (PPIs) in AtMad2 homocomplexes. We found that AtMad2 can be organized into dimers and tetramers (Fig. 4; Supplementary Fig. 10 and Supplementary Table 6). The phosphorylation of S190 blocks Mad2 dimerization (Kim et al., 2010), and BubR1 has been previously shown to block Mad2 dimerization in the MCC (Mariani et al., 2012). Furthermore, the interaction of Mad3 (BubR1 in yeast) with the mad2 dimer is functional for SAC progression (Mariani et al., 2012). We found that the interaction between AtMad2 (central region) and CDC20 (blade 3) is blocked by BubR1, more specifically by the conserved KET box motif (Fig. 3c–I,k,n; and Supplementary Tables 4–6). These results corroborate studies on SAC regulation via the interaction of BubR1 with the Mad2 homocomplex (Mapelli et al., 2007; Mariani et al., 2012; Faesen et al., 2017). Our findings suggest a regulatory mechanism in which AtBubR1 can block the interaction of the AtMad2 dimer (or tetramer) with AtCDC20, interfering with SAC signalling. The Mad2 dimer is formed by two antiparallel Mad2 molecules, where the hydrophobic core is important for the interaction among the open and closed structural conformations of these molecules (Mapelli et al., 2007). In this context, the formation of a tetramer can generate a structural arrangement with a central hydrophobic core delimited by four R134 residues (MIM of AtMad2). The importance of hydrophobic amino acids for interactions with the CDC20 MIM and the conversion of Mad2 from the open state to the closed state has been demonstrated recently (Yu et al., 2025). Some of these residues, L13, F23 and F24 (L14, F24 and F25 in AtMad2), are part of the hydrophobic core of Mad2 (Mapelli et al., 2007). Thus, we can speculate that the hydrophobic core in the AtMad2 tetramer may combine with hydrophobic interactions and hydrogen binding to trigger structural rearrangement, enabling autocatalytic conversion of O-Mad2 into C-Mad2 in the absence of Mad1 or Cdc20. However, a structural model of the AtMad2 tetramer is needed to analyse the implications of the assembly of four AtMad2 MIM motifs within this homocomplex, thereby confirming or refuting the hypothesis of an autocatalytic mechanism. In this context, Mad2 tetramers arrest cell cycle progression *in vitro* by inhibiting APC/C (Fang et al., 1998), and residues R133 and Q134 are required for the interaction between BubR1 and the Mad2 dimer. The presence of B9 and B10 crosslinks (Fig. 4e–h; Supplementary Table 5) suggests that the C-terminus of AtBubR1 can stabilize AtMad2 oligomers (i.e., dimers or tetramers) at the core MCC, supporting the hypothesis that the Mad2 dimer is a component of the MCC (Mariani et al., 2012). However, whether BubR1 can stabilize Mad2 tetramers in the MCC in *A. thaliana* and prevent cell cycle progression by impairing MCC-APC/C (Fang et al., 1998) still needs to be investigated in plants. Additionally, the MIM motif sequence of AtMad2 is predicted to interact with 14-3-3 and casein kinases (Fig. 4i). In plants, these two kinases are related to cell cycle regulation, tissue development, stress responses, metabolism regulation, and growth control (Youn et al., 2015, Zhao et al., 2021). Mutations in R134 analogues are sufficient to disrupt the SAC (Sironi et al., 2001; Mapelli et al., 2007; Fang et al., 1998; Hara et al., 2015). We speculate that phosphorylation at this specific point has the potential to disturb SAC progression. This could connect cell cycle control to variations in the immediate environment. In yeast and human cells, the core MCC adopts the same general structure, forming a heterotrimer with CDC20, Mad2 and BubR1 (Kops et al., 2020; Musacchio 2015; Musacchio and Salmon, 2007; McAinsh and Kops 2023). Here, we used full-length proteins and showed that more than one AtCDC20 can interact with AtBubR1, together with AtMad2 homocomplexes, to form the core MCC (Figs. 2, 3, and 5; Supplementary Figs. 11-14 and Supplementary Tables 4-6). In this context, we identified not only canonical PPIs but also novel plant-conserved and nonconserved motif binding. For instance, “motfis” KEN boxes are crucial for MCC training and SAC functions (King et al., 2007 and Tipton et al., 2011). With respect to AtBubR1 (BMF2), we identified three KEN boxes and one noncanonical KET box (Fig. 3e–g, l–n; Fig. 6f, Supplementary Fig. 5, Supplementary Table 5). We also revealed that there are shared interaction sites among AtMad2, AtCDC20 and AtBubR1, suggesting a complex mechanism of reciprocal regulation (Supplementary Figures 10-11). The identification of two functional binding sites for CDC20 on BubR1 with implications for the functionality of the SAC is not recent (Davenport et al., 2006). In addition, sequential coordination of the interaction between Mad2, BubR1 and CDC20 was suggested, where Mad2 interacts with CDC20, and the latter is then able to bind to BubR1 (Davenport et al., 2006). The binding affinities of AtMad2, AtCDc20 and AtBubR1 differ. The KIM motif of AtMad2 interacts preferentially with AtBubR1 in the KET box motif (Fig. 3e–g). The interaction of the KIM motif with the β-propeller domain occurs only in the absence of AtBubR1, which binds to the same region of AtCDC20. These findings suggest that AtBubR1 may regulate the interaction between AtMad2 and AtCDC20. These findings indicate that higher concentrations of AtMad2 and AtCDC20 are required for reciprocal interactions in a scenario in which the AtBubR1 concentration remains constant. In this context, the interaction of AtBubR1 with two AtCDC20 molecules in the absence of APC/C suggests that the stoichiometric balance of AtCDC20 is crucial for the functionality of the core MCC in *A. thaliana*. Corroborating this hypothesis, AtBubR1 can interact with itself (Supplementary Figure 13), suggesting that the C-terminus of AtBubR1 can “close” in a coiled-coil structure without AtCDC20 being available. The structural and functional characteristics of the CDC20 protein, its interactions with SAC proteins (BubR1 and Mad2) and its implications for SAC signalling have been extensively studied (Luo et al., 2000; Sironi et al., 2001; Tian et al., 2012; Hardwick et al., 2000; Fang et al., 1998; Davenport et al., 2006; Izawa et al., 2012; Tang et al., 2001; King et al., 2007; Yu et al., 2025, Luo et al., 2002). However, the identification of conserved MIM3 prompted us to investigate structural elements that enable the interaction of AtCDC20’s MIM3 with AtMad2 (Fig. 2e, f; Supplementary Figure 3). In this context, several examples of transient localized flexibility within the β-propeller domain as a requirement for specific functions exist. For instance, studies involving G proteins have shown that interactions with phosducin induce state changes in the β-propeller domain, specifically in blades 6 and 7 (Loew et al., 1998). More recently, it was shown that this type of change in the β-propeller domain can be reversed (Gatsogiannis et al., 2018). The absence of “Velcro” closure between the β-sheets of blades 1 and 7 allows for a certain degree of flexibility and enables access to the interior of the β-propeller domain on prolyl oligopeptidase (Fulop et al., 1998). In the β-sheets that interact between blades 1 and 7 of AtCDC20, strong hydrophobic interactions are unlikely, indicating a loosening of the “velcro” in this region. (Fig. 6a, b). Interestingly, we identified a conserved glycine-rich sequence (glycine box) that, in the context of the SAC, can form hydrogen bonds with KEN box 1 (AtBubR1) and the Mad2 dimer interface (Fig. 6c–e). We speculate that the interactions of the glycine box (AtCDC20) with both AtBubR1 and Mad2 homocomplexes may facilitate a transient state of disarray in blades 6 and 7. This could cause a slight displacement of blade 6, creating space for separation between the β-sheets of blades 1 and 7 (Fig. 6 a–e; Supplementary Figs. 3 and 4). In this way, an additional MIM motif could be exposed, resulting in CDC20 acting as another template for the conversion of O-Mad2 to C-Mad2. Acting as a “catalytic step” outside kinetochores (Musacchio 2015). This mechanism has the potential to accelerate the availability of C-Mad2 and may shift the kinetic balance towards the completion of SAC signalling. Our findings corroborate those of previous studies that revealed that the formation of Mad2 dimers accelerates the binding of Mad2 and CDC20 (Simonetta et al., 2009). We recognize the need for a specific structural model of the core MCC in *A. thaliana* to overcome limitations regarding further details on the interactions between AtCDC20, AtBubR1 and AtMad2. Taken together, our results show that in *A. thaliana*, AtBubR1 can function as a platform that accommodates more than one AtCDC20. In this context, AtCDC20 itself, in turn, also acts as a platform to accommodate homocomplexes (dimers or tetramers) of AtMad2 (Fig 7c). This reveals a significant difference in the composition of the core MCC compared with what is known in other eukaryotic models. In conclusion, despite structural similarities to those of other eukaryotes, we also identified interactions that are novel and specific to *A. thaliana*. While laying the structural foundation for core MCC formation in plants, our results also offer new insights into core MCC assembly in other eukaryotes.

## Methods

### Protein expression and purification

The full-length sequences of the following genes optimized for insect cells were obtained (GeneScript, New Jersey, EUA): AtCDC20 (At4g33270), AtBubR1 (AT2g33560), AtMad2 (At3g25980), AtBub1 (AT2g20635) and AtBub3;1 (AT3g19590). These genes were subsequently cloned and inserted into the pLIB vector (Weissmann et al., 2016). AtCDC20 was tagged with maltose-binding protein (MBP), and all the other proteins were tagged with 6His. To generate the plant core MCC, the following components were combined in pBIG1a (Weissmann et al., 2016): pLIB MBP AtCDC20, pLIB 6his AtBubR1 and pLIB AtMad2 6his. Additionally, individual pLIB 6his AtBub1 and pLIB 6his Bub3;1 constructs were produced. Baculoviruses were generated in SF9 insect cells and expressed in a 250 ml BTI-Tna038 insect cell suspension (Trowitzsch et al., 2010) in SF-900 III 1× medium (Gibco). After 96 h, these cells were pooled by centrifugation (4,000 rpm, 5 min, 4 °C). Afterwards, the supernatant was discarded, and the cells were resuspended in buffer containing 50 mM HEPES (pH 7.5), 300 mM NaCl, 5% glycerol, 10 mM imidazole and 2 mM TCEP (B300i), lysed by sonication and cleared by centrifugation (78,800 × g, 30 min and 4 °C). After the sediment was discarded, the supernatant was subjected to affinity chromatography (1 h, 4 °C and constant agitation) with a complete His-tag (Sigma‒Aldrich) according to the manufacturer’s instructions utilizing B300i buffer. The electrophoretic profile of this sample was analysed via 12% tricine‒ SDS‒PAGE and stained with Coomassie brilliant blue. Afterwards, the eluted sample was concentrated (Amicon tube 10 kDa), aliquoted, frozen in liquid nitrogen and stored at −80 °C.

### Size-exclusion chromatography

SEC was performed in a Superose 6, 10/300 GL column (GE Healthcare) equilibrated in 25 mM HEPES buffer (pH 7.5), 50 mM NaCl, 5% glycerol and 2 mM TCEP (B50) according to the recommended flow rates and isocratic conditions in a cold room (4 °C). After analyses of the chromatogram and electrophoretic profile obtained via 12% tricine-SDS‒PAGE with Coomassie brilliant blue staining, the fractions of interest were pooled, concentrated (Amicon tube 10 kDa), frozen in liquid nitrogen and stored at −80 °C.

### Pull-down assay using amylose resin

A pull-down assay was performed as described by Pan and collaborators (Pan et al., 2019), with a few modifications. The proteins of the plant core MCC, either alone or together with AtBub1 or Atbub3;1, were diluted to a concentration of 6 µM in 40 µl of 20 mM HEPES buffer (pH 7.5) containing 50 mM NaCl, 5% glycerol and 2 mM TCEP (B50) and gently mixed with 25 µl of amylose resin (NEB) that was previously equilibrated with buffer B50. Afterwards, 20 µl of this mixture was taken as the input sample, and the remaining sample (40 µl) was incubated for 30 min at 4 °C with constant agitation. The mixture was subsequently washed by centrifugation with 1 ml of buffer B50 three times to separate the amylose-bound proteins from the unbound fraction. After the proteins were eluted from the amylose resin by incubation with buffer B50 plus 10 mM maltose (Sigma‒Aldrich), they were incubated for 1 h at 4 °C with constant agitation. Next, the amylose was separated by centrifugation and discarded. The supernatant (eluted) was recovered and analysed via tricine‒SDS‒PAGE (12%) with Coomassie brilliant blue staining.

### Chemical crosslinking with DSBU

A chemical crosslinking assay was performed as described by Pan and collaborators (2018) with a few modifications. Proteins of the plant core MCC complex at a concentration of 3 µM (0.34 mg/ml) were dissolved in buffer containing 20 mM HEPES (pH 7.5), 50 mM NaCl, 5% glycerol and 2 mM TCEP (B50). Additionally, the plant MCC complex (2 µM (2.4 mg/ml)) plus 1 µM (1.2 mg/ml) AtBub1 or 1 µM (1.4 mg/ml) AtBub3:1 was dissolved in the same buffer (B50). Before the crosslinking reaction began, a 10 µl aliquot was collected for electrophoretic analysis. DSBU (Alinda Chemical Limited) was added to the protein solution at a final concentration of 3 mM. The samples were incubated at 25 °C for 1 h. The reaction was then stopped by the addition of Tris-HCl (pH 8.0) at a final concentration of 100 mM, followed by incubation at 25 °C for 30 min. A 10 µl aliquot was subsequently collected for analysis via tricine-SDS‒PAGE (12%) with Coomassie blue staining.

### Mass spectrometry analysis

Mass spectrometry analysis was performed as described by Hubner and collaborators (Hubner et al., 2010) with a few modifications. The samples were subsequently centrifuged at 15,000 rpm for 5 min, after which the supernatant was removed. Each sample was subsequently subjected to LysC/trypsin digestion, denaturation in 25 μL of denaturation–reduction buffer (8 M urea + 1 mM DTT) and vortexing at 25 °C for 30 min. The sample was subsequently subjected to alkylation with 5.5 mM chloroacetamide, followed by 20 min of vortexing at 25 °C. Next, 25 μL of 20 mM ammonium bicarbonate (ABC) buffer was added to achieve a final urea concentration of < 4 M, after which 2 μg of LysC was added, and the mixture was vortexed for 3 h at 25 °C (0.5 μg/μl LysC, 4 μl). Afterwards, 1 µg of trypsin (10 μl stock solution of 0.1 μg/μl + 40 μl dilution buffer) was added, and the sample was incubated at 25 °C overnight in dilution buffer (100 mM Tris, 1 mM CaCl2, pH 8.5). The next day, the digestion was stopped, with a final TFA concentration of 0.2%. The pH was approximately 2.5, and the final volume of the mixture was approximately 100 μL. The samples were then subjected to LC MS/MS analysis. MaxQuant searches to check the purity of the complexes were performed in parallel in a bacterial database, an insect database, and among the taxonomies of *Trichoplusia ni* and *Arabidopsis thaliana*. The sequences were submitted to a database of known contaminants with a false discovery rate (FDR) of 1%. The MeroX output was filtered to the experimentally observed mass deviation as indicated in the data file name, and the minimum score was set according to an FDR of 5%. Graphical representations showing intra- and intermolecular nonredundant crosslinks in circular plots were generated using xVis (https://xvis.genzentrum.lmu.de/CrossVisNoLogin.php).

### Computational tools

Sequence alingments were performed with MUSCLE with defaults in jalview v 2.11.5.1 (Waterhouse et al., 2009). Sequence logo charts were generated with a local installation of weblogo v3 (Crooks et al., 2004). Protein structure models were generated using Modeller protocols (Webb & Sali, 2017). Cartoon representations of protein structures and measurements between Cα–Cα with nonredundant crosslinks were prepared using PyMOL (available at https://pymol.org/2) following the instructions in the user guide. To predict protein domains and short linear motifs (SLiMs), we used ELM (Kumar, et al., 2020). Cartoons for the topologic representation of protein domains and different motifs were generated using Domain Graph version 1.0 (DOG) (Ren et al., 2009).

## Acknowledgements

Thank you to Andrea Musacchio for inviting me and giving me the opportunity to achieve this work. We are very grateful for working at the Max Planck Institute in Dortmund. Special thanks to Dongqing Pan for his excellent tips and relevant discussions. We also thank Sabine Wohlgemuth, Isabelle Stender, Dorothee Vogt, Carolin Koerne and Franziska Müller for their technical support. Thanks to Gilliana Dutra Neves Cruz for her comments and suggestions in the preparation of this manuscript. We thank the Max Planck Institute of Molecular Physiology and all of Department 1. This work was supported by Max Planck Society - Max-Planck-Gesellschaft (127493).

## Ethics

### Author contributions

E.A.G.O. and E.V.N.G. collected the data and performed the models; I.S.S. collected the data and performed the computational analysis; G. P. S and P.G. C. They carried out data collection and performed alignments, consensus analyses, and sequence topologies; M.A.L.C. conceived the project, performed protein expression and purification, designed the protein experiments, and prepared and wrote the manuscript.

### Data availability

