## Supplementary information (figures and tables) for "Core MCC of *Arabidopsis thaliana*: structural dissection provides new insights into Mad2, BubR1 and CDC20"

### Supporting information

### Supplementary Figure 1

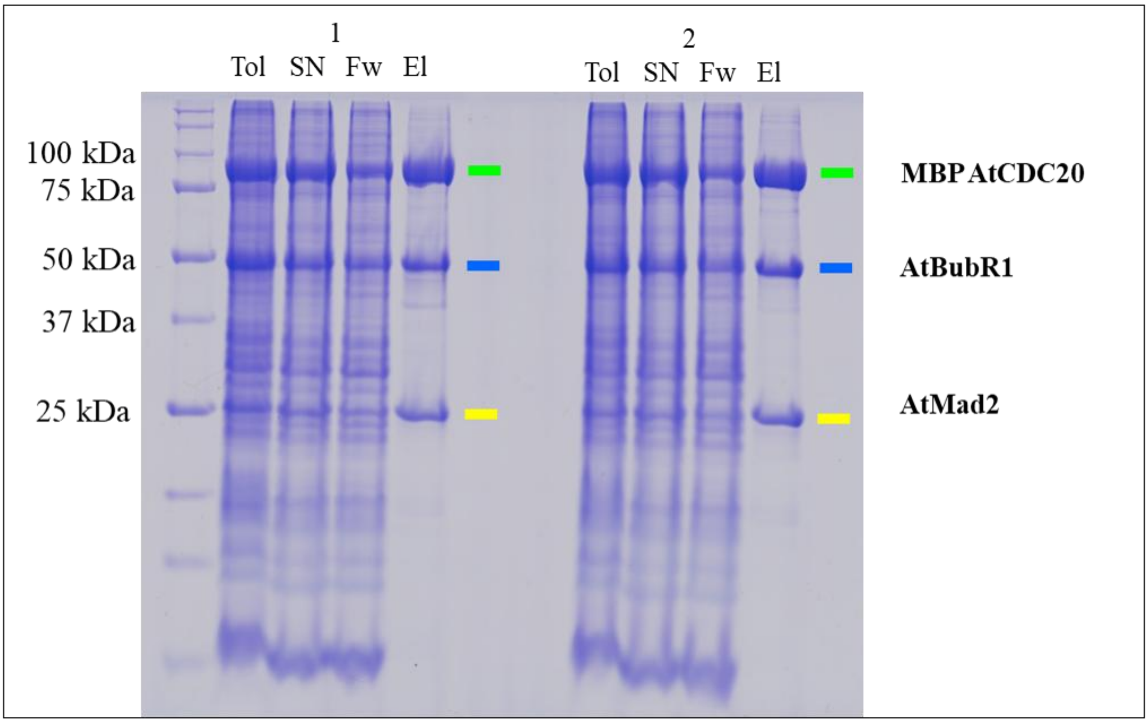


**Supplementary Figure 1 Expression of the bigBac core MCC proteins of *Arabidopsis thaliana* a**, SDS–PAGE expression analysis of the core MCC protein of *Arabidopsis thaliana* after the first step of purification using affinity chromatography. M – marker, Tol- total lysate, SN- supernatant, Fw- discarded flow through, El – eluate.

**Supplementary Figure 2**

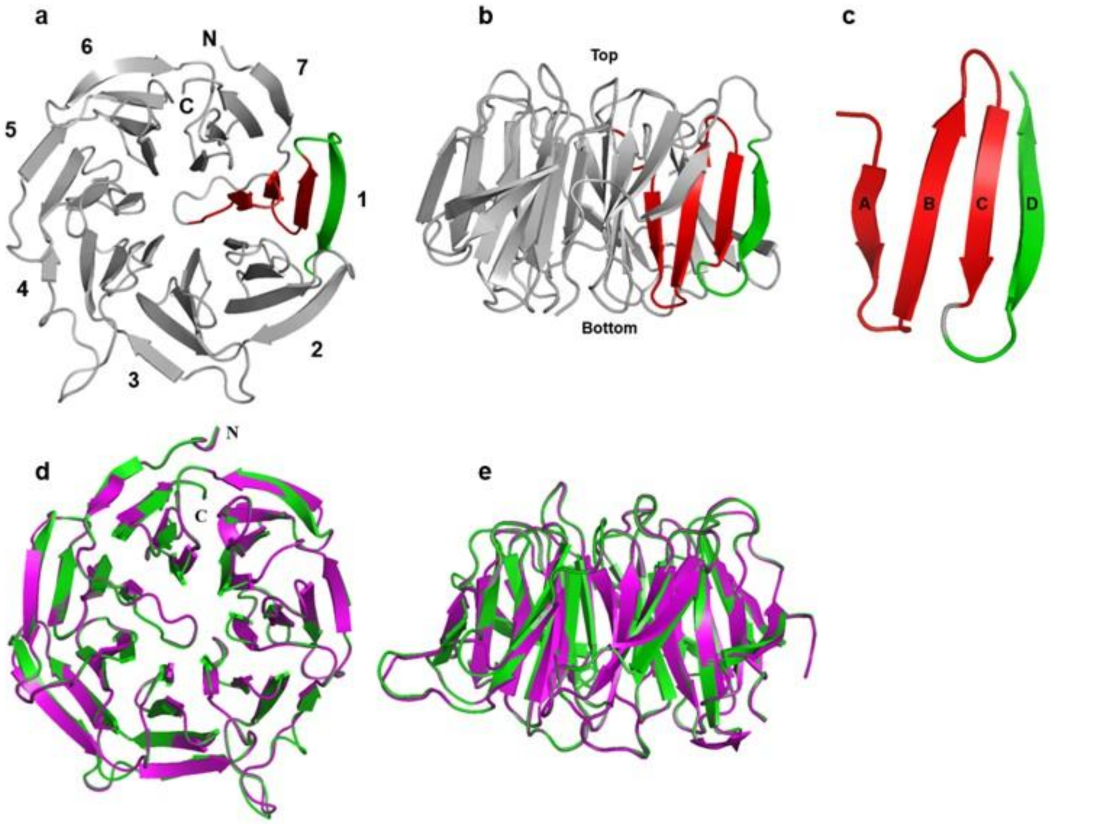


**Supplementary Figure 2 β-propeller domain of AtCDC20 a, b**, Blade organization of the β-propeller domain of AtCDC20 with WD40 repeats (blades). **c**, Details of the blade with a β-sheet organization, where each blade is formed by four antiparallel β-sheets, including three blades that are the same (A, B and C) (red) and the β-sheet (D) of the next blade (green). **c**, **d**, Top view of the superposition of the β-propeller domains of AtCDC20 (green) and HsCDC20 (magenta). **e**, Lateral view of the superposition of the β-propeller domain.

**Supplementary Figure 3**

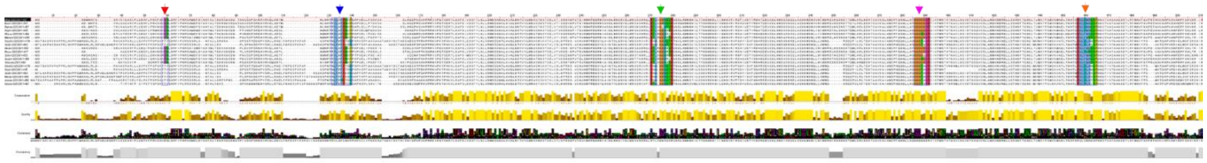

**Supplementary Figure 3 - Plant CDC20 alignment -** Red arrows indicate the KEN box motif at the N-terminus. The blue arrows indicate the MIM1 motif sequence. The green arrows indicate the MIM2 motif sequence. Pink arrows indicate the glycine box sequence. The orange arrows indicate the MIM3 sequence.

**Supplementary Figure 4**

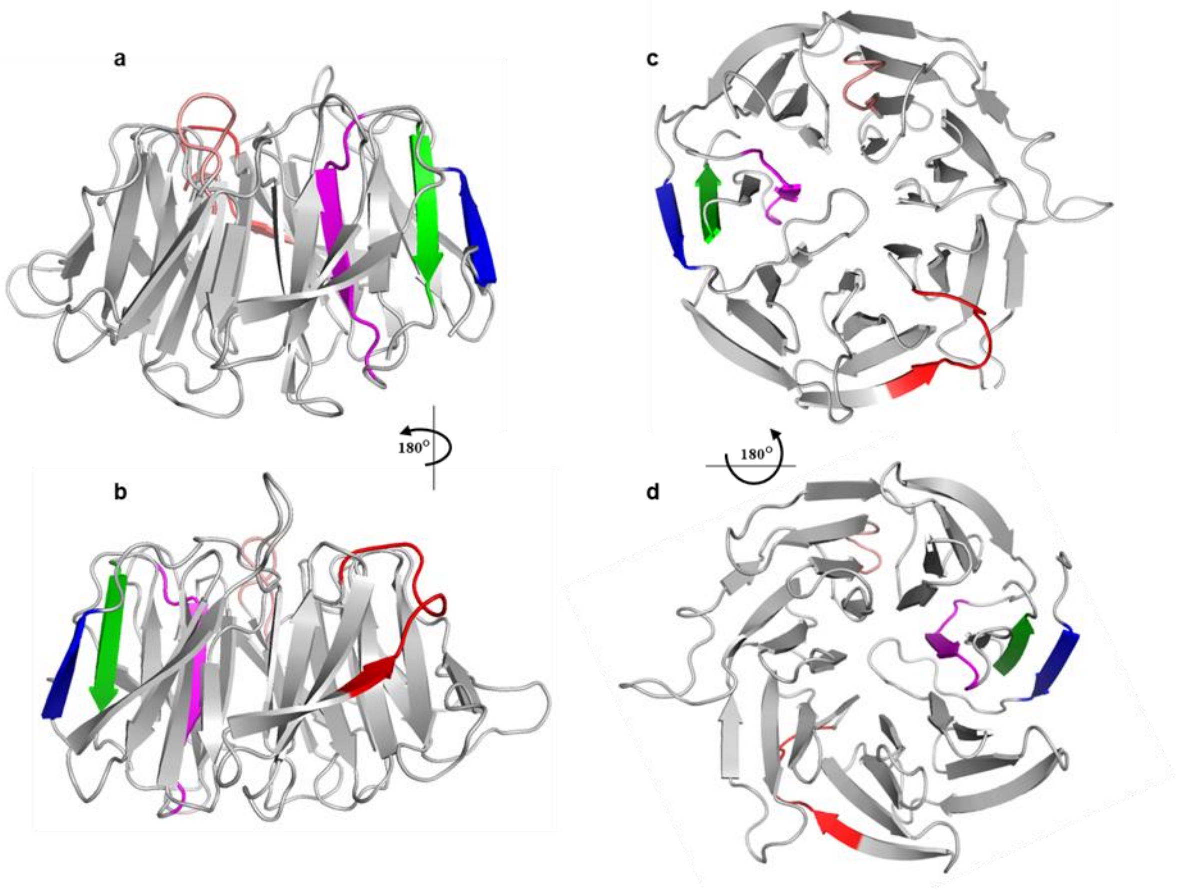


**Supplementary Figure 4 - Structural characteristics of the β-propeller domain of AtCDC20 a,** Lateral view of the β-propeller domain showing the structural characteristics of AtCDC20 in grey. MIM2 (red), MIM3 (magenta), glycine box (salmon), β-sheet of blade1 (blue) and β-sheet of blade 7 (green) of AtCDC20. The same colour scheme was used throughout the figure. **b**, Lateral view of the β-propeller domain showing structural characteristics with 180° rotation. **c**, Top view of the β-proporter domains of AtCDC20. **d**, Bottom view of the β-propeller domains of AtCDC20.

**Supplementary Figure 5**

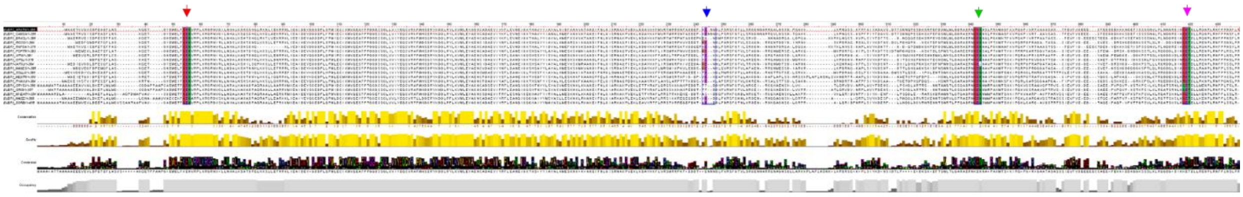

**Supplementary Figure 5 - Plant BubR1 alignment -** Red arrows indicate the KEN box_1 motif at the N-terminus. The blue arrows indicate the KEN box_2 sequence. The green arrows indicate the KEN box_3 sequence. The blue arrows indicate the KET box sequence.

**Supplementary Figure 6**

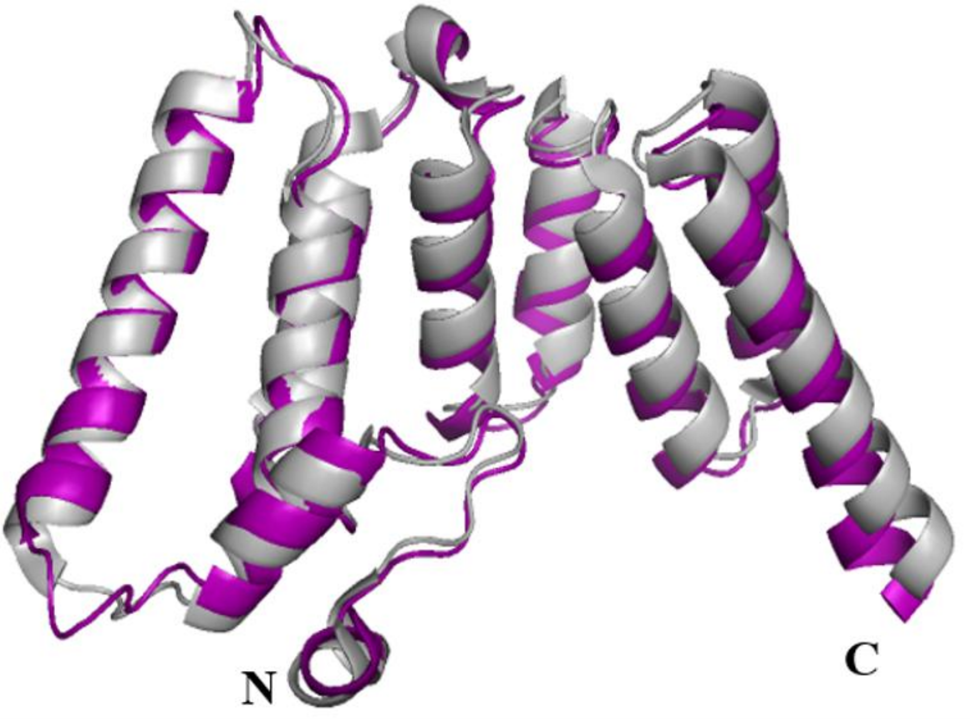


**Supplementary Figure 6 BubR1 TPR domain -** Superposition of the N-terminus and TPR domain with AtBubR1 (purple) and HsBubR1 (gray). N- and C-terminal indicate

**Supplementary Figure 7**

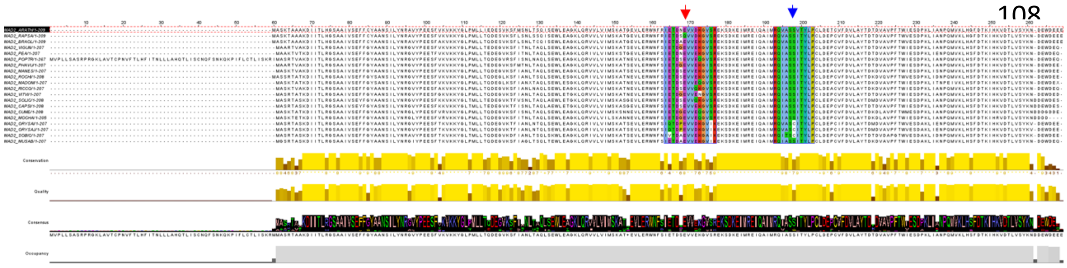

**Supplementary Figure 7 - Plant Mad2 alignment -** Red arrows indicate the KIM motif of AtMad2. The blue arrows indicate the AtMad2 MIM sequence.

**Supplementary Figure 8**

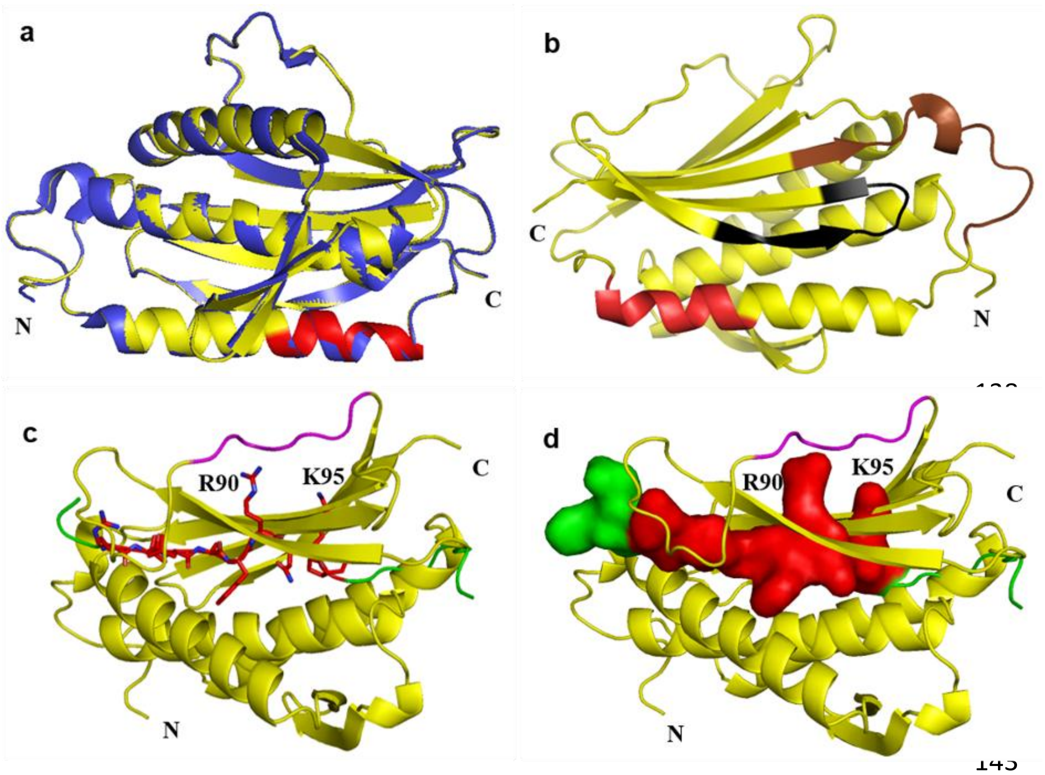


**Supplementary Figure 8 Supplementary Figure 4 General structure of AtMad2 a,** Cartoon representation showing the superposition of C-AtMad2 (yellow) and C-HsMad2 (blue) and the AtMad2 MIM motif (red). **b**, Cartoon representation of the AtMad2 model showing the protein–protein interaction position in the AtMad2 MIM motif (red) and hotspots in the central region (brown) and the C-terminal region (black). **c** and **d**, Cartoons showing the AtCDC20 MIM motif in red (sticks and surface) near the “safety belt” (purple) in the C-terminus of AtMad2 (yellow). AtCDC20 (green) and the amino acids arginine and lysine (90 and 95, respectively) in the AtCDC20 MIM motif are indicated in black.

**Supplementary Figure 9**

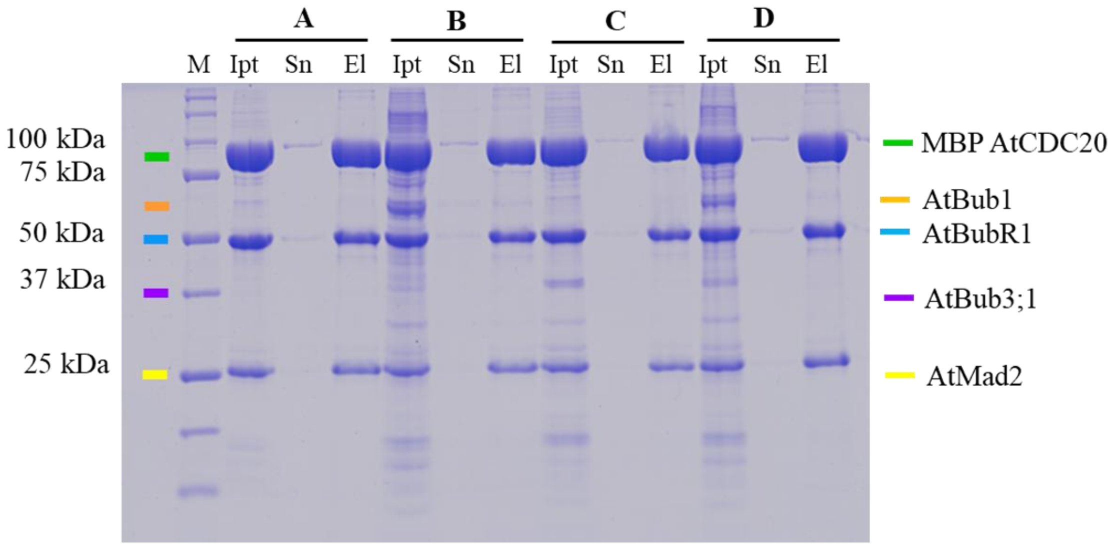


**Supplementary Figure 9 - Pull-down assay using amylose resin -** SDS–PAGE analysis of the results of pull-down with amylose and MBP AtCDC20 used as bait: sample A: core MCC (MBP AtCDC20, 6his AtBubR1 and 6his AtMad2), sample B: core MCC plus 6his AtBub1), sample C: core MCC plus 6his AtBub3;1, sample D: core MCC plus 6his AtBub1 and 6his AtBub3; M: marker; Ipt: input; Sn: supernatant and El: eluted.

**Supplementary Figure 10**

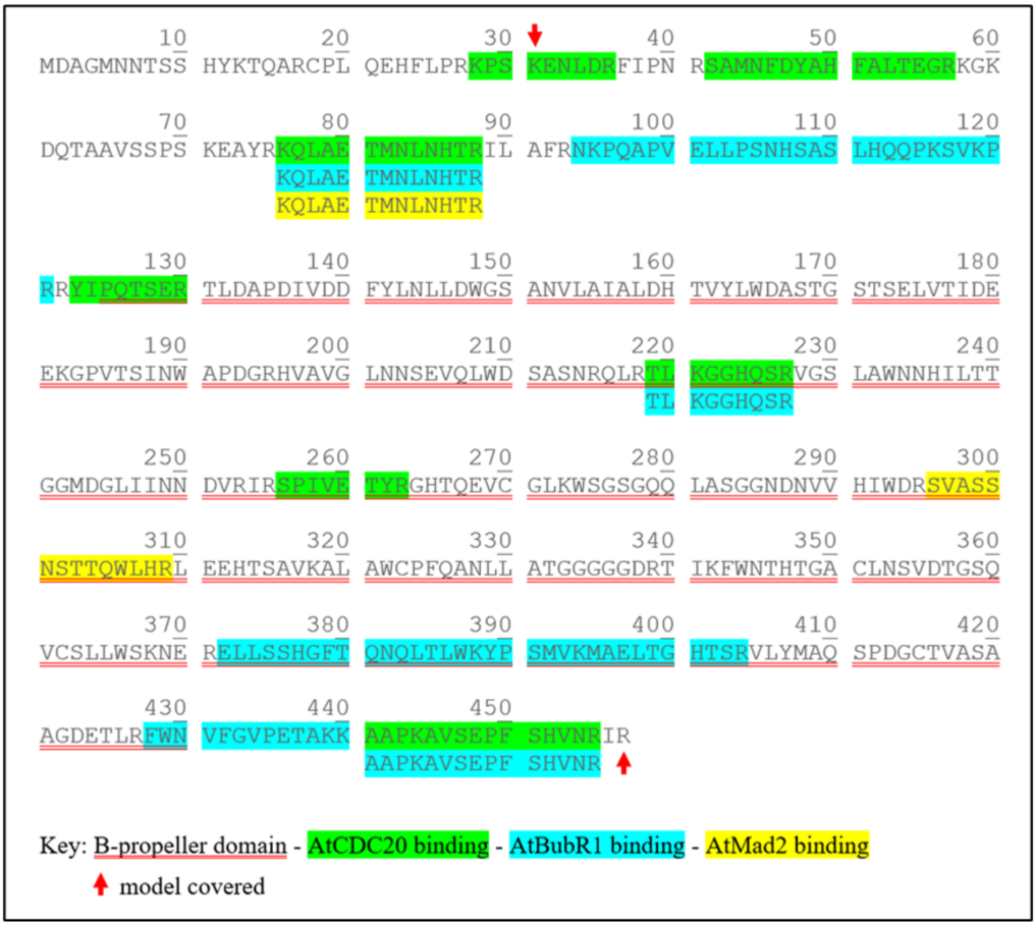


**Supplementary Figure 10 - Mapping of the nonredundant crosslinking of AtCDC20 -** Sequence showing protein–protein interaction sites identified via chemical crosslinking with DSBU.

**Supplementary Figure 11**

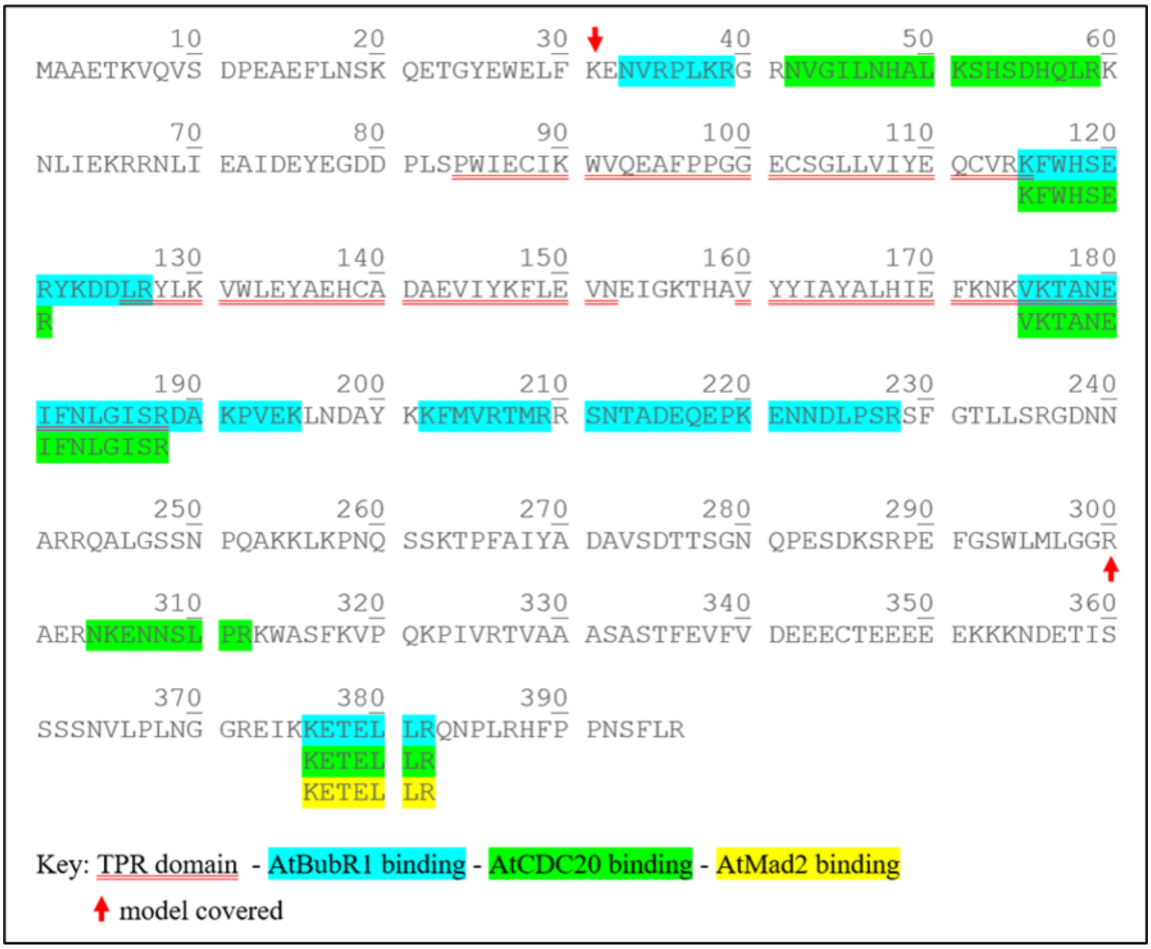


**Supplementary Figure 11 - Mapping of the on-redundant crosslinking of AtBubR1 -** Sequence showing protein–protein interaction sites identified via chemical crosslinking with DSBU.

**Supplementary Figure 12**

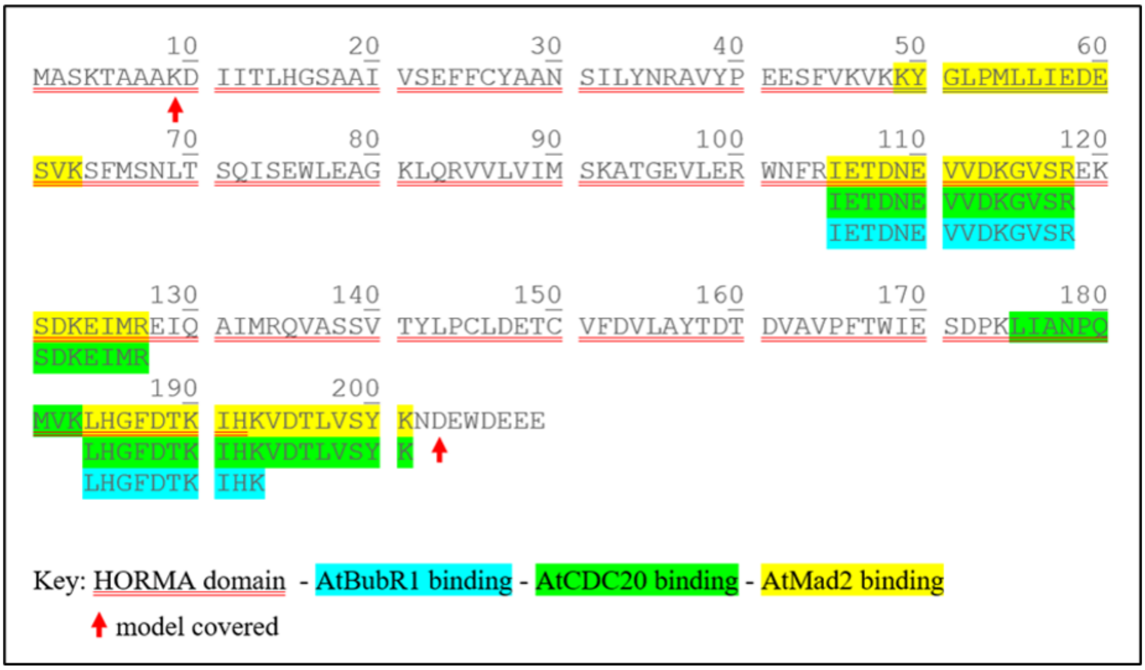


**Supplementary Figure 12 - Nonredundant crosslinking of AtMad2**; sequence showing protein–protein interaction sites identified via chemical crosslinking with DSBU.

**Supplementary Figure 13**

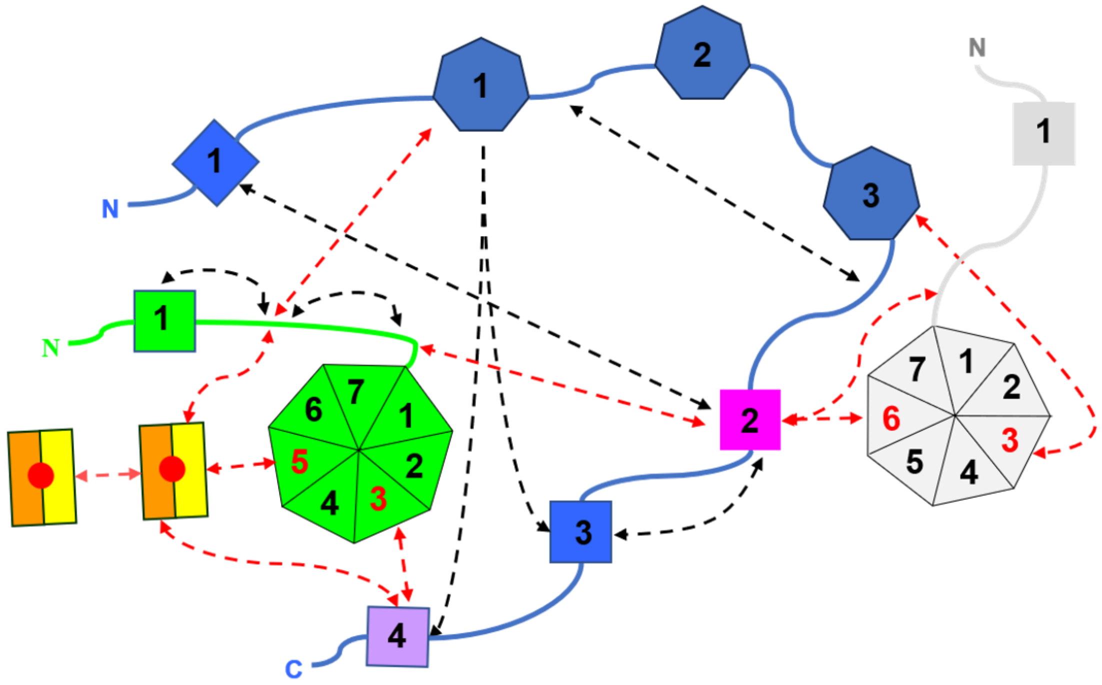


**Supplementary Figure 13 - Protein–protein interactions of the core MCC in *Arabidopsis thaliana* -** Interaction mapping scheme of protein–protein interactions between AtBubR1, AtCDC20 and AtMad2 according to the chemical nonredundant crosslinking results. The colour scheme is indicated in the key. The black dashed arrows indicate interactions mapped in the models used. Dashed red arrows indicate at least one peptide outside the models used.

**Supplementary Figure 14**

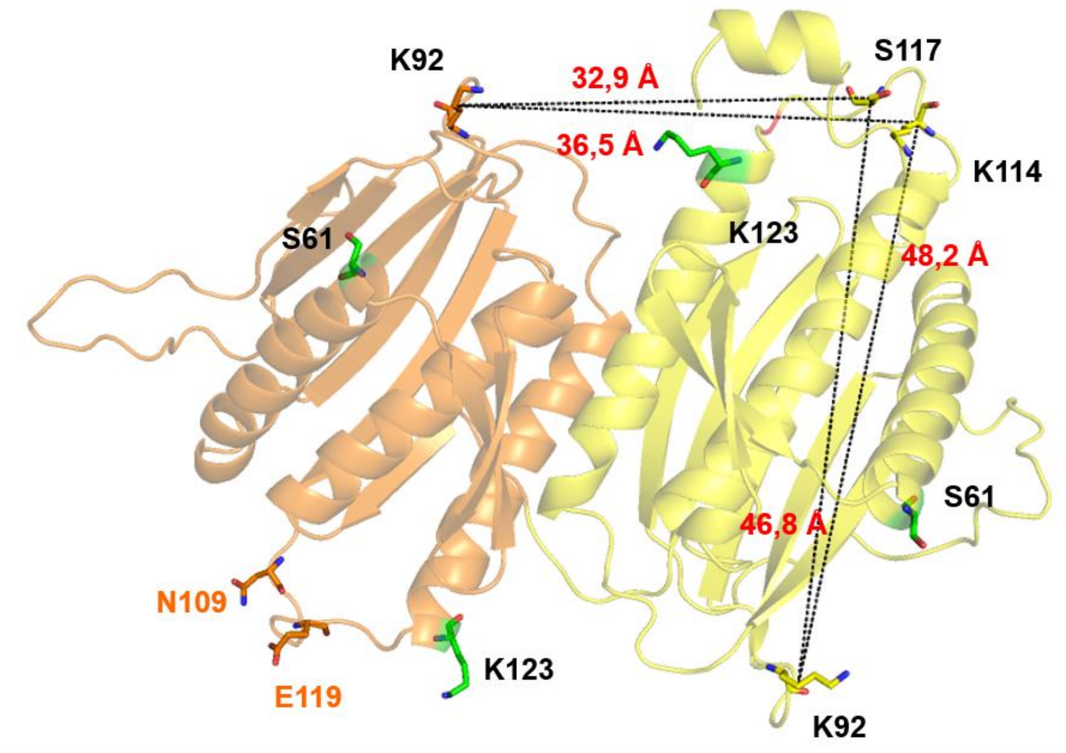


**Supplementary Figure 14 - Distances between the amino acids involved in the nonredundant crosslink M9 and M10 -** Cartoon representation of the Mad2 dimer; the distances in angstroms (red) between amino acids K92 and K114 or S117 are shown as dashed black lines. O-Mad2 (orange) and C-Mad2 (yellow). Colours: nitrogen (blue), oxygen (red). N109 and E110 (orange) are shown as a reference for hypothetical positions K114 and S117. Amino acids involved in the nonredundant crosslink M8 (green). Model 2v64 was used (Mapelli et al., 2007).

**Supplementary Table 1** – Model parameters of the core MCC of *Arabidopsis thaliana.* Template (Alfieri et al., 2017)**.**

| Component | Template | Chain | Start position (aa) | Final position (aa) | Coverage (%) |
|---|---|---|---|---|---|
| AtCDC20$^{mc1}$ | 5LCW | "Q" | 85 | 457 | 372 |
| AtBubR1mcc | 5LCW | "S" | 24 | 369 | 345 |
| AtMad2mcc | 5LCW | "Z" | 9 | 204 | 195 |
| AtCDC20$^{mc2}$ | 5LCW | "R" | 31 | 457 | 426 |

**Supplementary Table 2** – Model parameters of the mitotic complex of *Arabidopsis thaliana.* Template (Chao et al., 2012)**.**

| Componente | Template | Chain | Start position (aa) | Final position (aa) | Coverage (%) |
|---|---|---|---|---|---|
| AtCDC20y$^{mc1}$ | 4aez | "A" | 85 | 431 | 75,7 |
| AtBubR1ymcc | 4aez | "C" | 26 | 248 | 62,2 |
| AtMad2ymcc | 4aez | "B" | 15 | 201 | 96,2 |

**Supplementary Table 3** – Model parameters of the AtMad2 dimer of *Arabidopsis thaliana.* Template (Mapelli et al., 2007)**.**

| Component | Template | Chain | Start position (aa) | End position (aa) | Total (%) |
|---|---|---|---|---|---|
| Mad2's dimmer | 2v64 | "A" | 15 | 206 | 84,2 |
| Mad2's dimmer | 2v64 | "D" | 10 | 206 | 86,6 |

**Supplementary Table 4 Nonredundant crosslinking of AtCDC20** – Sequence showing protein–protein interaction sites identified via chemical crosslinking with DSBU.

| Type | Cross-links | Residue 1 (AtCDC20) | Residue 2 (Protein) | Distance Cα- Cα (Å) | Times observed | Cross-link experiment |
|---|---|---|---|---|---|---|
| Intramolecular | C1 | T54 | AtCDC20 K76 | 18.1 | 92 | D |
| | C2 | K76 | AtCDC20 K31 | -- | 35 | D |
| | C3 | K76 | AtCDC20 Y123 | 22.1 | 24 | D |
| | C4 | K76 | AtCDC20 Y123 | | 18 | E |
| | C5 | K221 | AtCDC20 S226 | 10.2 | 6 | A |
| | C6 | K221 | AtCDC20 S256 | 10.4 | 88/104/132 | A/C/D |
| | C7 | K221 | AtCDC20S 254 | 11.1 | 9 | E |
| | C8 | K388 | AtCDC20 S451 | 30.6 | 10 | D |
| Intermolecular | C9 | K76 | AtBubR1 S54 | -- | 10 | D |
| | C10 | K76 | AtBubR1 K115 | -- | 110 | D |
| | C11 | K76 | AtBubR1 K119 | -- | 43 | A |
| | C12 | S110 | AtBubR1 K176 | 29.1 | 12 | D |
| | C13 | S110 | AtBubR1 K220 | 18.2 | 9/22/67 | A/C/D |
| | C14 | K119 | AtBubR1 K176 | 29.0 | 10 | D |
| | C15 | K119 | AtBubR1 S211 | 28.1 | 56/33/106/147 | A/B/C/D |
| | C16 | K119 | AtBubR1 S227 | 20.5 | 9/26/39 | A/C/D |
| | C17 | K221 | AtBubR1 K176 | 14.2 | 66/111 | A/D |
| | C18 | K221 | AtBubR1 K376 | -- | 152/85 | A/C |
| | C19 | S226 | AtBubR1 K376 | -- | 164/74 | C/D |
| | C20 | T380 | AtBubR1 K220 | 35.7 | 50 | D |
| | C21 | K388 | AtBubR1 K220 | 54.1 | 29 | D |
| | C22 | K388 | AtBubR1 S227 | 26.3 | 65/7/172 | A/C/D |
| | C23 | K394 | AtBubR1 S227 | 22.3 | 131 | D |
| | C24 | K439 | AtBubR1 S227 | -- | 15 | D |
| | C25 | K444 | AtBubR1 S227 | -- | 44 | D |
| | C26 | K71 | AtMad2 S117 | -- | 129 | D |
| | C27 | K76 | AtMad2 S117 | -- | 126/130/139 | A/C/D |
| | C28 | K76 | AtMad2 S189 | -- | 79 | D |
| | C29 | K76 | AtMad2 K190 | -- | 64 | D |
| | C30 | K76 | AtMad2 K193 | -- | 110 | D |
| | C31 | T81 | AtMad2 S183 | -- | 44 | D |
| | C32 | T81 | AtMad2 K190 | -- | 43 | D |
| | C33 | T87 | AtMad2 K114 | 17.4 | 89 | A |
| | C34 | K76 | AtMad2 K183 | -- | 21 | E |
| | C35 | K76 | AtMad2 K189 | -- | 18 | E |
| | C36 | K76 | AtMad2 K193 | -- | 8 | E |
| | C37 | T81 | AtMad2 K114 | -- | 148 | E |
| | C38 | K221 | AtMad2 K183 | -- | 9 | E |
| | C39 | K76 | AtMad2 S117 | -- | 43 | E |
| | C40 | S300 | AtMad2 K123 | 18.6 | 100 | D |
| | C41 | T303 | AtMad2 K123 | 16.9 | 91 | D |
| | C42 | K221 | AtMad2 S117 | -- | 29 | E |
| | C43 | K221 | AtMad2 K114 | -- | 6 | E |

**Supplementary Table 5** – Summary of the nonredundant crosslinking of AtBubR1 in experiments A, B, C and D

| Type | Cross-link | Residue 1 (AtBubR1) | Residue 2 (Protein) | Distance Cα-Cα in (Å) | Times observed | Cross-link experiments |
|---|---|---|---|---|---|---|
| Intramolecular | B1 | K38 | AtBubR1 K202 | 18.1 | 84/85 | A/D |
| | B2 | K115 | AtBubR1 K305 | -- | 106 | D |
| | B3 | K115 | AtBubR1 K376 | -- | 128/146 | A/D |
| | B4 | K123 | AtBubR1 K191 | 14.0 | 44/157 | A/ C |
| | B5 | K176 | AtBubR1 S211 | 13.5 | 68 | A |
| | B6 | S227 | AtBubR1 K176 | 14.1 | 122 | A |
| | B7 | S227 | AtBubR1 K305 | -- | 66/110/87 | A/C/D |
| Intermolecular | B8 | T376 | AtMad2 K190 | -- | 46 | D |
| | B9 | T378 | AtMad2 K114 | -- | 85/91 | A/D |

**Supplementary Table 6** – Sequence showing protein–protein interaction sites identified via chemical crosslinking with DSBU.

| Type | Cross-link | Residue 1 (AtMad2) | Residue 2 | Distance (Cα-Cα in Å) | Times observed | Cross-link experiment |
|---|---|---|---|---|---|---|
| Intramolecular | M1 | K114 | AtMad2 S117 | 6.6 | 180/9/188 | A/C/D |
| | M2 | K114 | AtMad2 K190 | 12.2 | 119/57 | D/E |
| | M3 | S117 | AtMad2 K123 | 15.1 | 52/50 | D/E |
| | M4 | S117 | AtMad2 K190 | 10.1 | 57/106/77/84 | A/C/D/E |
| | M5 | S117 | AtMad2 K193 | 16.2 | 25 | D |
| | M6 | K190 | AtMad2 S121 | 10.5 | 51/8 | A/C |
| | M7 | K190 | AtMad2 K123 | 13.4 | 79/25/142 | A/C/D |
| Intermolecular | M8 | AtMad2 S61 | AtMad2 K123 | -- | 8/10 | A/D |
| | M9 | AtMad2 K92 | AtMad2 K114 | -- | 8 | E |
| | M10 | AtMad2 K92 | AtMad2 S117 | -- | 22 | E |
