## Supplementary figures and images for "Core MCC of *Arabidopsis thaliana*: structural dissection provides new insights into Mad2, BubR1 and CDC20"

### Supplementary figure 3

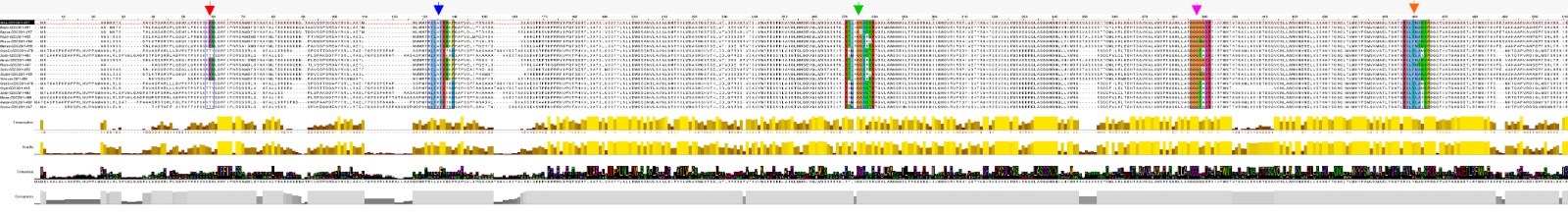

### Supplementary figure 5

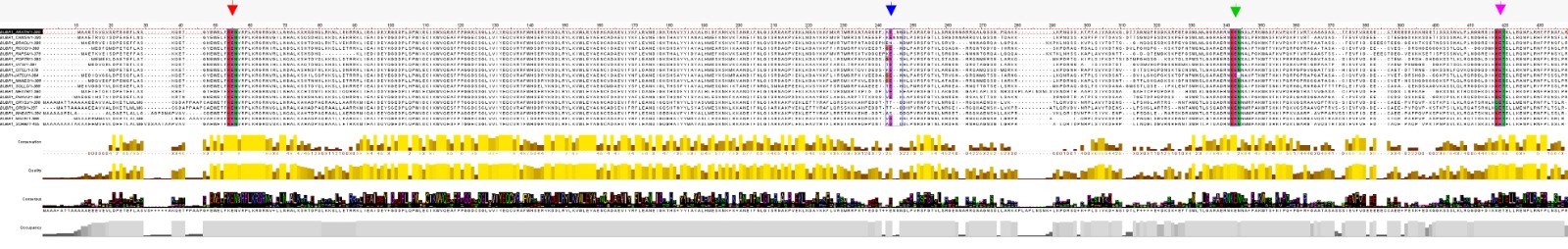

### Supplementary figure 7

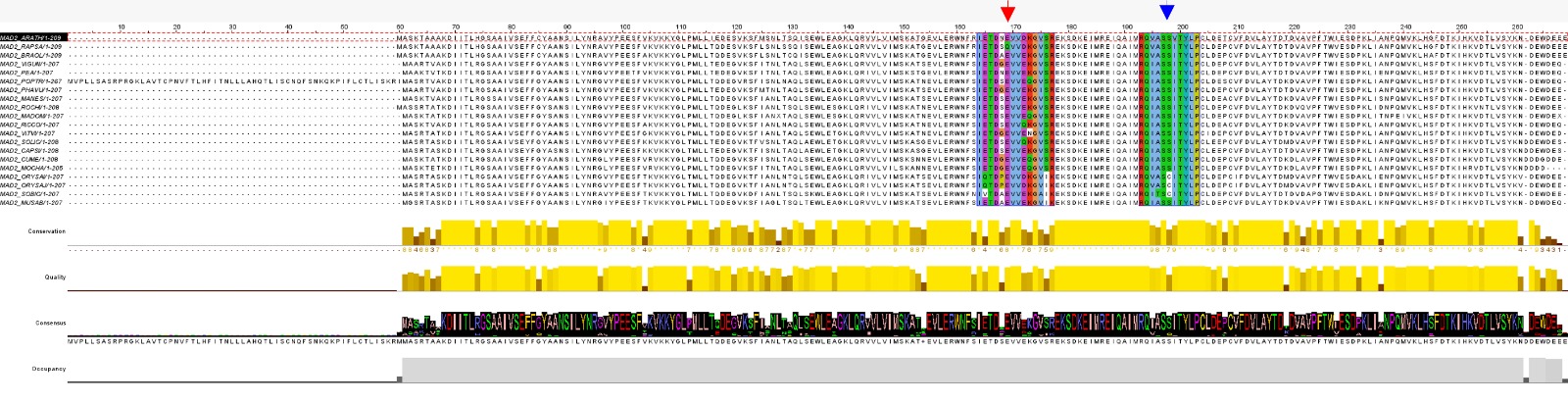
